# Holobiont transcriptomes for the critically endangered staghorn coral (*Acropora cervicornis)* from two environmentally distinct sites on Turneffe Atoll, Belize

**DOI:** 10.1101/2022.03.29.486305

**Authors:** Kathryn C. Lesneski, Adam T Labadorf, Karina Scavo Lord, Filisia Agus, John R. Finnerty

**Affiliations:** Boston University Biology Department, 5 Cummington Mall, Boston, MA 02215; Marine Program, 5 Cummington Mall, Boston, MA 02215; Boston University School of Medicine, 75 Concord Street, Boston MA 02129

**Keywords:** holobiont, transcriptome, *Acropora cervicornis*, Belize

## Abstract

Historically, staghorn coral (*Acropora cervicornis*) was a preeminent reef-builder in the Caribbean and Tropical Western Atlantic, where it constructed extensive thickets at 5-20 m depth that supported diverse ecosystems and provided coastal populations with food, storm protection, and income from tourism. In recent decades, *A. cervicornis* declined precipitously, up to 97% in some localities. To reverse its decline, widespread efforts are underway to characterize the phenotypic and genetic diversity of persisting populations with the goal of restoring them to historical levels by out-planting nursery grown specimens. To support this target, we developed transcriptomes for two *A. cervicornis* populations located in Turneffe Atoll Marine Reserve, Belize. These populations experience significantly different temperatures, light levels and water currents, and they harbor individuals that differ in key phenotypes. Because differentiating the gene activity of diverse taxa—i.e., coral host, algal photosymbiont, plus associated eukaryotes, bacteria, archaea, and viruses— is critical to understanding the function of the coral holobiont, we developed a pipeline for parsing transcripts by taxon. Separate transcriptomes for each population contain complete representatives for >96% of 978 conserved metazoan single copy orthologs. The taxonomic breakdown of transcripts differed between sites, with more bacterial transcripts recovered from Calabash Caye and more symbiont transcripts from Blackbird Caye. The assembled transcriptomes will facilitate gene expression studies and *in silico* cloning from this endangered coral.

## Introduction

Historically, staghorn coral, *Acropora cervicornis* (Lamarck 1816), was a principal reef builder across the Caribbean and Tropical Western Atlantic (TWA), typically the dominant species at 5-25 m on exposed forereefs, and widespread in backreef and lagoonal habitats [1]. Since the 1970s, staghorn coral has undergone a precipitous decline, with some areas experiencing up to a 97% loss [2]. The formerly dense cover of *A. cervicornis* provided habitat for fishes and invertebrates, supporting the fishing and tourism industries. It also attenuated wave energy, reducing the impact of coastal storms and enabling the establishment of seagrass and mangrove ecosystems [3].

Given its former prominence and ecological importance, *A. cervicornis* is a central focus of reef restoration efforts across the Caribbean and TWA. Thus, understanding the phenotypic and genetic background of corals reared in nurseries and outplanted onto reefs is key to long-term success of restoration efforts. Genomic and transcriptomic data can aid in identifying resilient genets and revealing the molecular mechanisms underlying differences in resilience [4].

Here we describe reference transcriptomes for two *A. cervicornis* populations from Turneffe Atoll Marine Reserve, Belize. The Blackbird Caye backreef and Calabash Caye forereef apron (i.e. backreef to forereef transition) differ significantly in key environmental parameters, but both harbor *A. cervicornis* colonies that have persisted over the last several years. We parsed annotated contigs into taxonomic bins for coral host, photosymbionts (Symbiodiniaceae), and ten other taxonomic classes, which will facilitate taxonomically focused analyses of gene expression. Given the high level of genetic diversity across the Caribbean [5], the existence of high-quality transcriptomes for the western Caribbean, Belize in particular, will inform genetically based conservation efforts. The taxonomic parsing method we describe should be useful for annotating any holobiont transcriptome.

## Materials and Methods

### Site selection and environmental characterization

Corals were collected near Blackbird Caye (BC) and Calabash Caye (CC), mangrove islands located 4.0 km apart on the eastern side of Turneffe Atoll, Belize (Fig. 1). Surveys in 2014 revealed scattered *A. cervicornis* colonies in the backreef at BC and along the forereef apron at CC, at depths of approximately 1.0 m and 2.5-3.0 m, respectively. From 2014-2018, we recorded temperature at each site using Onset® HOBO® Pendant data loggers every 3-6 h. The same loggers were used to measure light intensity during the first 6-7 days of deployment, before fouling of the sensors by encrusting organisms. Water current (m/s) was recorded every 10 minutes during a 70-hour period in November 2019 using Marotte HS current meters (Marine Geophysics Laboratory, Townsville, Australia).

**Figure 1.**
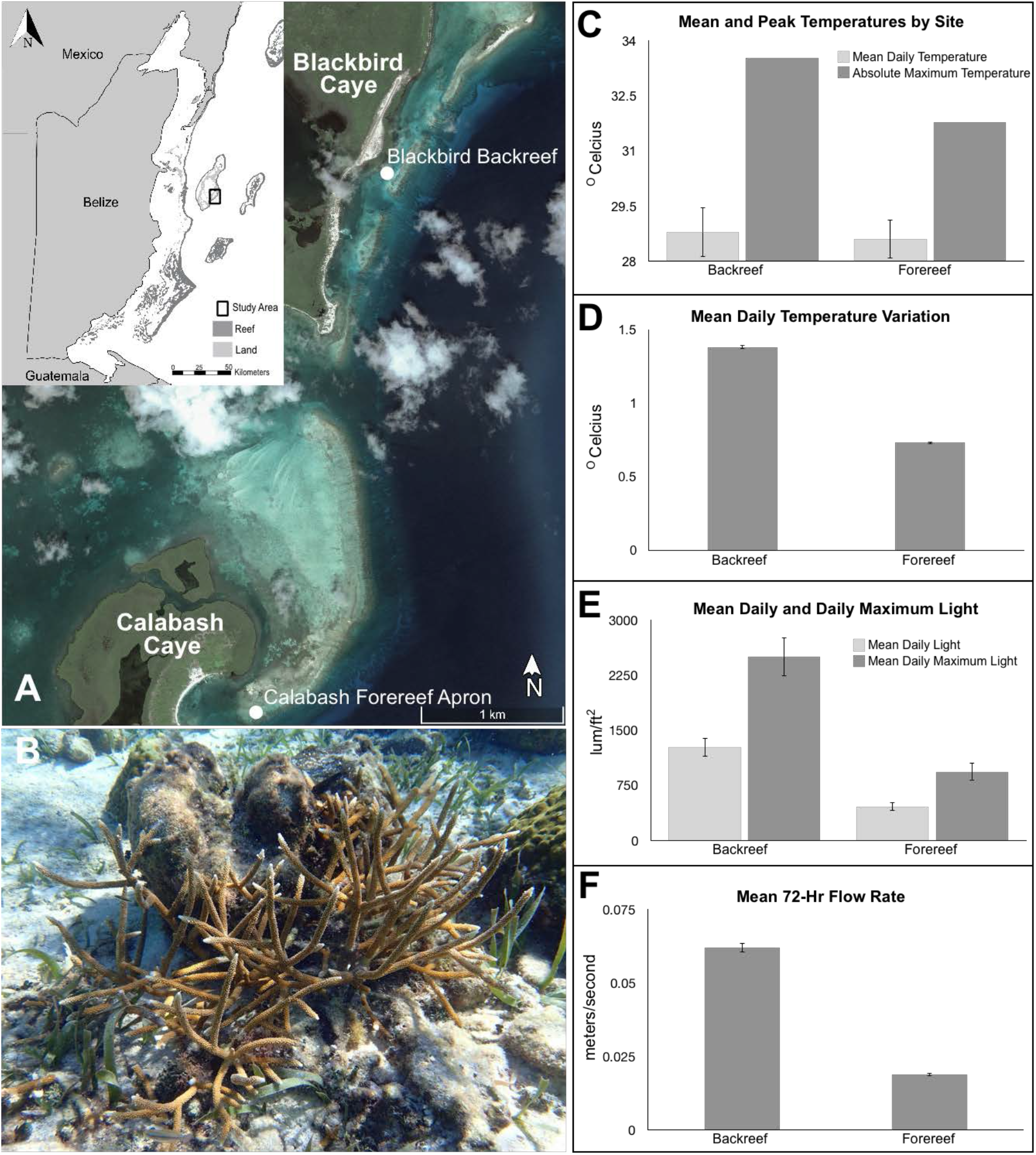
Study site and environmental data Collection sites and environmental monitoring. **(A)** Blackbird Caye and Calabash Caye, Turneffe Atoll, Belize (Google Earth Pro, v 7.3.2.5776; image date: August 26, 2005, © 2020 Maxar Technologies). **(B)** *Acropora cervicornis* colony at Blackbird Caye. **(C)** Four-year averages for mean and peak temperature by site. **(D)** Mean daily temperature variation. **(E)** Light intensity measured over four one-week periods during each study year. **(F)** Mean flow rate by site, measured over a 72-hour period.

### Sample collection and RNA extraction

In November 2017, during early morning (0600-0730), a ∼1 cm branch tip was collected from each of four spatially isolated *A. cervicornis* colonies at Blackbird Caye (BC) and Calabash Caye (CC) and placed in 100% ethanol. Within 1 h, the ethanol was replaced with fresh ethanol, and samples were stored at -20 °C. Sample metadata are provided in Table 1. Approximately 0.1 cm^3^ of skeletal tissue was crushed using a razor blade and combined with acid-washed 425-600 µm glass beads in Lysis/Binding solution from the RNAqueous™ Total RNA Isolation Kit (ThermoFisher Scientific). The resulting mixture was agitated in a Fisherbrand™ Bead Mill 24 Homogenizer for 1 minute to remove tissue from skeletal fragments. Total RNA from four BC and four CC colonies was extracted using the RNAqueous™ kit and sent to Novogene for sequencing.

**Table 1.** Specimen metadata

|  | **Blackbird** | **Calabash** |
| --- | --- | --- |
| Organism Part | Branch tip (n=4; 1 per colony) | Branch tip (n=4; 1 per colony) |
| Lifestage | Adult | Adult |
| Project Name | Blackbird Caye *Acropora cervicornis* transcriptome | Calabash Caye *Acropora cervicornis* transcriptome |
| Collected By | Kathryn C. Lesneski | Kathryn C. Lesneski |
| Collection Date | November 2017 | November 2017 |
| Country | Belize | Belize |
| Latitude | 17.314444 N | 17.28 N |
| Longitude | 87.797222 W | 87.807778 W |
| Locality, Region | Blackbird Caye, Turneffe Atoll | Calabash Caye, Turneffe Atoll |
| Habitat | Back reef | Fore reef apron |
| Sex | Hermaphrodite | Hermaphrodite |
| Collecting Institution | Boston University | Boston University |

### Sequencing and de novo assembly

Data generation and analysis are summarized in Suppl. Fig. 1. Libraries were created by Novogene using the NEBNext® Ultra™ DNA LibraryPrep Kit (New England Biolabs) and sequenced on an Illumina Novaseq6000 using paired-end, 150-bp reads. Adaptors were removed by Novogene. Raw fastq files were assessed using FastQC [6]. Sequencing reads were trimmed and filtered with Cutadapt [7]. The first 15 nt were trimmed from forward and reverse reads, because they exhibited a biased composition. Both members of a read pair were discarded if either read exhibited phred score q<30 or length <20 nt.

We generated multi-kmer assemblies for BC and CC using Velvet-Oases [8, 9]. We produced single-kmer assemblies at 55, 65, 75, 85, 95, 105, and 115 nt, and then merged at the default kmer (27 nt). Each assembly was assessed for completeness using BUSCO (version 3.0.2; [10]), to compare transcripts generated in this study with the metazoa_odb9 and Alveolata datasets. N50 values were obtained using TransRate [11]. Each transcriptome was mined for molecular markers that have been widely characterized in Symbiodiniaceae (actin, COI, ITS2, rad24; [12]) and Cnidaria (NK-κB; IKKα/β, IKKε/TBK1; [13]); the resulting transcripts were assessed for completeness and analyzed phylogenetically (Suppl. Figs. 2-14).

### Examining relatedness of coral samples

As clonal reproduction via fragmentation is common in *A. cervicornis* [14], we analyzed single nucleotide polymorphisms (SNPs) mined from the transcriptome to assess relatedness between colonies (Suppl. Fig. 15).

### Transcriptome annotation

Assembled transcripts from both sites were concatenated into a single holobiont transcriptome. Sequences were clustered using CD-HIT [15] if they were at least 30% of each other’s length and shared 90% sequence identity. Transcripts were aligned against the NCBI Nucleotide (nt) database (accessed September 7, 2019) using *minimap2* [16]. Transcripts that failed to align were subsequently aligned against the NCBI Protein (nr) database (accessed September 7, 2019) using DIAMOND [17], with hits required to exhibit at least 50% coverage of the query. The match exhibiting the lowest e-value was chosen as the best hit. To break ties, we excluded protein names that contained “PREDICTED”, “LOW QUALITY PROTEIN”, or “hypothetical protein,” and then randomly chose one of the remaining best hits. Each transcript was assigned to one of 11 taxonomic classes based upon the taxon ID associated with its best nucleotide or protein match (Suppl. Table 1). If a transcript matched equally well to different IDs, the most general taxclass was chosen. Transcripts with no hit were mapped to the taxclass associated with the longest transcript in its CD-Hit cluster. If the longest transcript in the cluster also had no hit, the transcript was assigned to a “No Hit” bin.

### Taxonomic variation within and between sites

To identify unique and overlapping transcripts and genes in the holobiont transcriptomes by taxclass, we used *minimap2* [16] to align the Blackbird and Calabash transcriptomes. To quantify reads from each taxclass in each of the eight sequencing samples, the sequencing reads were aligned to the transcriptome of their parent site using *samtools* [18]. We developed a graph-based algorithm to visualize differences in taxonomic representation between sites according to three different properties: taxonomic purity, genic purity, and RNA copy number (i.e. expression level) purity (Suppl. Fig. 16-17).

To assess variation in taxonomic composition of the metatranscriptome between colonies, we mapped reads in the eight sequencing datasets to the best match among transcripts in the holobiont transcriptome using *salmon* [19]. The resulting read counts were concatenated using *csvgather* [20] and subdivided according to taxonomic class. All code to reproduce the taxonomic classification analysis is available on github at https://github.com/BU-Neuromics/staghorn_transcriptome. All taxgraph results, filtered counts, and fasta files from taxonomic parsing are available on OSF at https://osf.io/u9nq8/

### BLAST interface for site-specific and taxon-specific transcripts collections

Separate BLAST databases were created for the Blackbird and Calabash holobiont transcriptomes, and for the eleven taxon-specific transcript collections (Suppl. Table 1) using SequenceServer, v. 1.0.11 [21].

## Results

### Environmental Differences Between Collection Sites

From November 2014 – December 2018, the backreef at Blackbird Caye (BC) exhibited a higher average daily temperature (+ 0.18 °C), higher maximum temperature (+2.27 °C), and greater average daily temperature range (+ 0.65 °C) than the forereef apron at Calabash Caye (CC). During four week-long measurement periods in November of 2014-2017, mean daily and mean maximum light intensity (lum/ft^2^) averaged 3x greater at BC than CC. Over a 72-hour period in 2019, flow was significantly higher at BC (0.062 m/s) than CC (0.019 m/s) (Fig. 1).

### Sequencing Yield and Transcriptome Assembly Statistics

After quality filtering, sequencing of four BC and four CC libraries yielded 13.10 and 13.57 gigabases, respectively (NCBI Short Read Archive, accession PRJNA737066). The BC, CC, and combined holobiont assemblies comprise 586,964, 415,325, and 1,002,289 contigs, respectively, with N50 values of 3,093, 2,374, and 2,792 nt. All three holobiont transcriptomes, as well as the taxonomically-parsed transcript collections are available for download and blast [22]. Trimmed versions of site-specific assemblies were also submitted to the Transcriptome Shotgun Assembly archive at NCBI following removal of contigs shorter than 200 nt, contigs containing sequencing adaptors, and contigs flagged by NCBI as potential contaminants (accession number GJIF00000000).

The BC, CC, and combined holobiont transcriptomes compare favorably with other coral holobiont transcriptomes for recovery of conserved metazoan orthologs (Suppl. Fig. 2). Using BUSCO, we recovered complete sequences for 98.5%, 96.0%, and 98.6% of 978 metazoan orthologs from the BC, CC, and combined assemblies, respectively. Following taxonomic parsing, we recovered complete sequences for 98.2% of 978 metazoan orthologs from the collection of cnidarian transcripts. Targeted BLAST searches recovered complete transcripts of protein-coding genes previously studied in corals, including NF-κB signaling pathway members (Suppl. Table 2). A motif analysis using the MEME Suite [23] recovered all conserved motifs identified in a prior study of coral NF-κB signaling proteins [24], suggesting all transcripts were complete (Suppl. Figs. 10-14).

With respect to symbiont transcripts, we recovered a similar fraction of conserved alveolate orthologs from the BC and CC transcriptomes as observed in other corals [24-26] and a *Symbiodinium microadriaticum* transcriptome (Suppl. Fig. 2). Additionally, we recovered apparently full-length transcripts for *Symbiodiniaceae* actin, COI, ITS-2, and rad24 (GenBank MZ501686-MZ501696, MZ503278) from BC, though in some cases, we were only able to recover partial sequences from CC (Suppl. Table 3).

### Relatedness of Coral Samples

Based on 35,637 SNPs, all four CC samples appear to be clones, while three of four colonies collected at BC appear to be clones (Suppl. Fig. 15).

### Taxonomic Distribution of Transcripts and Genes

Taxonomic distribution of transcripts is similar across sites. Cnidarian transcripts predominate at BC (64.8%) and CC (68.8%), with the two sites sharing 96.3% of cnidarian transcripts (Fig. 2; Suppl. Table 4). Both transcriptomes exhibit a similar proportion of transcripts in taxclasses No Hit (14.4% vs. 16.6%), Other Metazoa (3.77% vs. 3.30%), Fungi (0.19% vs 0.14%), and Other Eukaryota (1.58% vs 0.67%). Transcripts from Archaea, Viruses, and Other Sequences amount to less than 0.01% of transcripts at both sites (Fig. 2; Suppl. Table 4).

**Figure 2.**
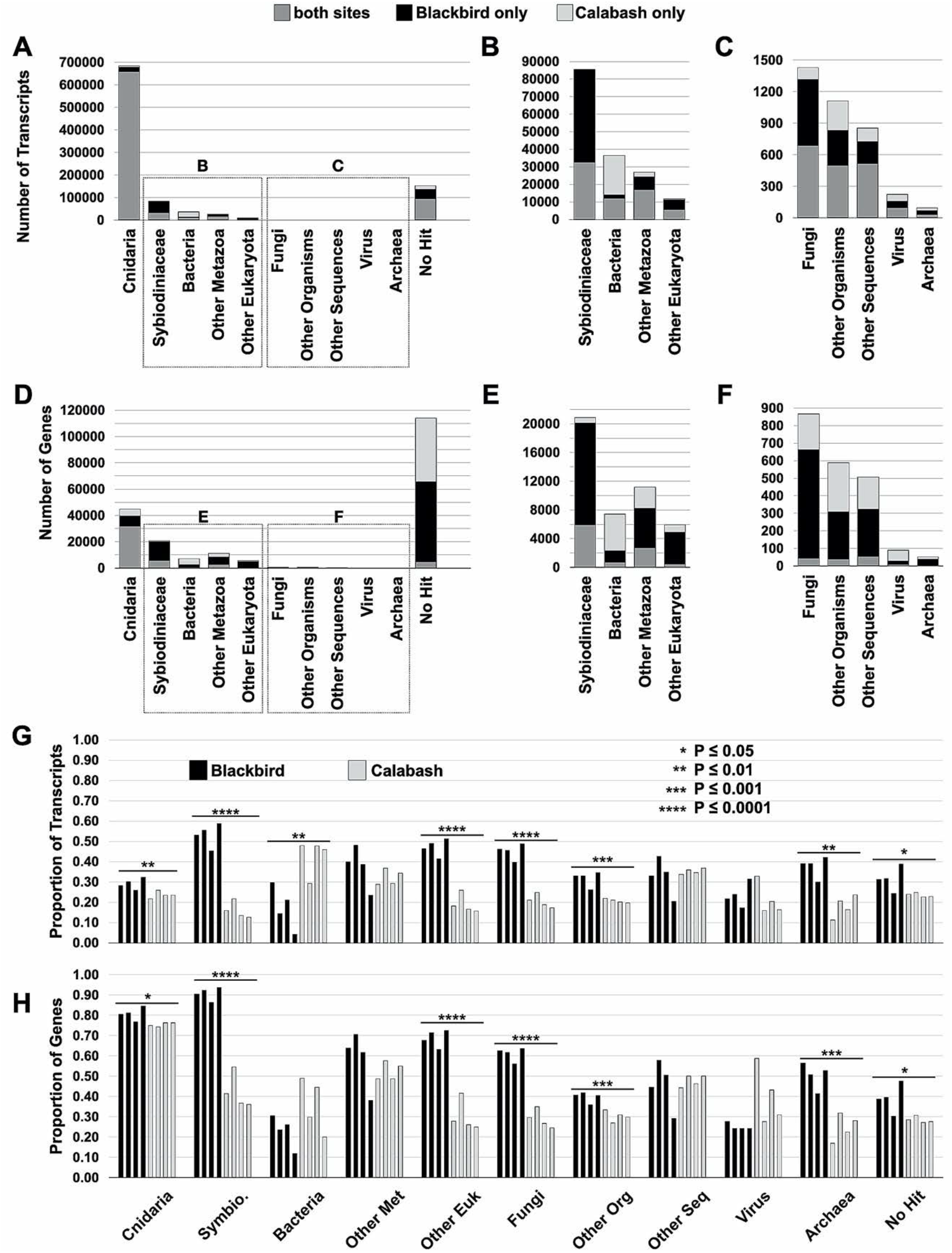
Variation across sites in the abundance of reads, transcripts, and genes by taxclass. Stacked bar charts for the number of transcripts **(A, B, C)** and number of genes **(D, E, F)** found only in Blackbird, only in Calabash, or at both sites for each of the 11 taxclasses. Proportion of transcripts **(G)** and genes **(H)** found in individual samples used in assembly of the separate Blackbird and Calabash transcriptomes. Significant differences (bars) and associated p-values (*) for the average total number of transcripts and genes between sites for each taxclass are represented here. T-tests or Kruskal-Wallis tests were conducted, depending on normality of data, to determine if the average total number of transcripts and genes significantly differed between sites for each taxclass for both transcripts and genes.

In contrast to other taxclasses, the proportion of transcripts from Symbiodiniaceae and Bacteria differ considerably between sites (Fig 2a-c). Symbiodiniaceae accounts for 13.9% of transcripts in BC but only 2.9% of transcripts in CC. The two sites share only 38.2% of transcripts in the taxclass Symbiodiniaceae (Suppl. Table 4). The taxclass Bacteria accounts for 1.0% of transcripts in BC but 7.2% in CC. Again, a relatively low fraction of transcripts is shared between sites (32.8%; Suppl. Table 4). For all taxclasses, a higher percent of genes compared to transcripts were unique to either site (Fig. 2d-f), implying that multiple transcripts are shared between sites for each gene that is shared between sites, on average.

To examine whether individual outliers are driving observed differences between sites, we plotted the average proportion of transcripts and genes for each individual sequencing dataset for each taxclass (Fig. 2g, h). Within sites, individual samples exhibit roughly equal proportions of transcripts and genes by taxclass. Between sites, the average number of transcripts (Fig. 2g) and the average proportion of genes (Fig. 2h) differs significantly for most taxclasses. The most highly significant differences between sites were the greater proportion of transcripts and genes from Symbiodiniaceae, Other Eukaryotes, and Fungi in BC.

## Discussion

The assemblies described here substantially augment the transcriptomic resources available for *A. cervicornis* in the Caribbean. Libro and co-workers (2013) developed a transcriptome based on combined RNA sequence data from *A. cervicornis* and *A. palmata* [27]. Combining species risks generating mosaic contigs. Furthermore, this transcriptome exhibited a relatively low N50 (363 nt), suggesting a paucity of full-length transcripts. By contrast, the site-specific assemblies generated here feature N50’s 6.5x (Calabash) and 8.5x (Blackbird) greater. Our ability to recover complete transcripts for >98% of conserved metazoan orthologs as well as select cnidarian (NF-κB, IKKα/β, IKKε) and Symbiodiniaceae genes (actin, COI, ITS-2, rad24) further indicates the assemblies are replete with full-length transcripts. Importantly, by separately assembling the data from each site, we facilitate the identification of site-specific differences in transcriptomic diversity and reduce the possibility of generating mosaic transcripts that don’t occur in either population. The ability to analyze transcriptomic variability at fine spatial scales may prove to be critical, as there is considerable population genetic structure in this species at distances of just 2 km [28]. Targeted conservation approaches based on the genetic profiles of individual populations may be necessary [5].

Numerous studies in the last 10-15 years have revealed the critical role the “holobiont” plays in the health of a coral [29]. Understanding the transcriptional activity of a holobiont requires identifying the taxonomic source of transcripts expressed under different environmental conditions or physiological states. Indeed, recent recommendations for coral restoration advocate a comparative approach involving holobionts sourced from various locations and spanning environmental gradients [30]. For this reason, the annotation pipeline we employed in this study incorporated a more exhaustive taxonomic parsing approach than has previously been used on any coral transcriptome.

Going forward, it will be important to identify factors (e.g., host genetic background or environment) driving differences in the metatranscriptome between sites. In the current study, we cannot disentangle the effects of environment and genotype on the observed differences between the metatranscriptomes of Blackbird and Calabash because different genotypes occupy different sites. To determine whether the observed differences in the diversity of bacterial and symbiont transcripts are driven by the environment or are stably associated with corals sourced from a particular location, we leveraged the transcriptomes described here to evaluate gene expression at 6 months, 1 year and 2 years in a reciprocal transplant experiment that we are currently preparing for publication (Lesneski et al., in prep).

## Author Contributions

**Kathryn C. Lesneski**: Conceptualization, Methodology, Formal analysis, Resources, Writing – Original Draft, Writing – Review & Editing, Visualization, Project administration, Funding acquisition. **Adam Labadorf**: Conceptualization, Methodology, Formal analysis, Resources, Writing – Original Draft, Writing – Review & Editing, Visualization. **Karina Scavo Lord**: Methodology, Resources, Writing – Review & Editing, Visualization. **Filisia Agus**: Methodology, Resources, Writing – Review & Editing. **John R. Finnerty**: Conceptualization, Methodology, Formal analysis, Writing – Original Draft, Writing – Review & Editing, Visualization, Project administration, Funding acquisition.

## Funding and Permits

The research described here was supported by National Science Foundation grant IOS-1354935 to JRF and by grants to KCL from: (1) the American Academy of Underwater Sciences (Doctoral Scholarship), (2) the American Philosophical Society (Field Research Grant), (3) the Explorer’s Club (Exploration Fund Grant), (4) the Flying Sharks Research Fund, (5) the Lewis and Clark Fund for Exploration and Research, (6) the Mohamed bin Zayed Species Conservation Fund, (7) Ocean Opportunity LLC, (8) the Robert & Patricia Switzer Foundation (Switzer Environmental Fellowship), (9) the Women Divers Hall of Fame, and (10) the Boston University Marine Program (Warren-McLeod Fellowship). The research was conducted at the Calabash Caye Field Station, Environmental Research Institute, University of Belize. Tissue sampling was conducted under Marine Scientific Research Permit #00055-17 (2017) and CITES Permit #6893 (2017) issued by the Belize Fisheries Department.

## Acknowledgements

We thank J. Azueta, I. Majil, and F. Cruz at Belize Fisheries Department for help obtaining research permits. Sample collection was performed by graduate students participating in Boston University’s Marine Semester. We thank the BU Marine Program, especially J. Hammer-Mendez, J. Perry, and J. Scace, for logistical and technical support, and S. W. Davies, B. Benson, and N. Kriefall for assistance with RNA extraction, C. Becker of MIT/WHOI for consultation on coral microbiomes and Jeffrey S. Sanborn of Boston University for deployment of the Blast database. This research required the local knowledge and technical expertise of boat captains, staff, and researchers at Calabash Caye Field Station and the University of Belize (M. Alamina, V. Alamina, A. Cherrington, L. Morey, L. Cho-Ricketts, N. Craig, M. Ipro). We thank the Bertarelli Foundation and the Oak Foundation for their contributions to the research infrastructure at Calabash Caye Field Station, which facilitated the marine conservation related research described here.

