## Supplemental Figures and Tables for "Holobiont transcriptomes for the critically endangered staghorn coral (*Acropora cervicornis)* from two environmentally distinct sites on Turneffe Atoll, Belize"

#### Supplemental Figures

| | | |
|---|---|---|
| **1** | Data generation and analysis pipeline | 2-3 |
| **2** | Recovery of universal, single-copy orthologs from cnidarian and Symbiodiniaceae transcriptomes | 4-5 |
| **3** | Alignment of Symbiodiniaceae actin proteins | 6-7 |
| **4** | Maximum likelihood phylogeny of Symbiodiniaceae actin | 8-9 |
| **5** | Alignment of Symbiodiniaceae COI proteins | 10 |
| **6** | Maximum likelihood phylogeny of Symbiodiniaceae cytochrome oxidase I (COI) | 11-12 |
| **7** | Maximum likelihood phylogeny of Symbiodiniaceae internally transcribed spacer 2 (ITS2) | 13-14 |
| **8** | Alignment of Symbiodiniaceae rad24 proteins | 15-16 |
| **9** | Maximum likelihood phylogeny of Symbiodiniaceae cytochrome rad24 | 17-18 |
| **10** | Alignment of coral and human NF-kB proteins | 19-21 |
| **11** | Conserved protein motifs in coral and human NF-kB proteins | 22 |
| **12** | Alignment of coral and human IKK a/b proteins | 23-25 |
| **13** | Alignment of coral and human IKKE/TBK1 proteins | 26-28 |
| **14** | Conserved protein motifs in coral and human IKK proteins | 29 |
| **15** | Acropora cervicornis SNP-based distance tree | 30-31 |
| **16** | Example calculations of taxon-, CPM-, and gene-purity scores | 32-33 |
| **17** | Graphs depicting the degree of taxonomic skewness between sites | 34-35 |

#### Supplemental Tables

| | | |
|---|---|---|
| **1** | Taxonomic class definitions and properties | 36 |
| **2** | Recovery of Symbiodiniaceae molecular markers from Blackbird and Calabash | 37 |
| **3** | Recovery of NFkB signaling components from Blackbird and Calabash | 38 |
| **4** | Number of transcripts shared by and unique to Blackbird and Calabash by taxclass | 39 |

#### Literature Cited 40

**Supplemental Figure 1**. Data generation and analysis pipeline.

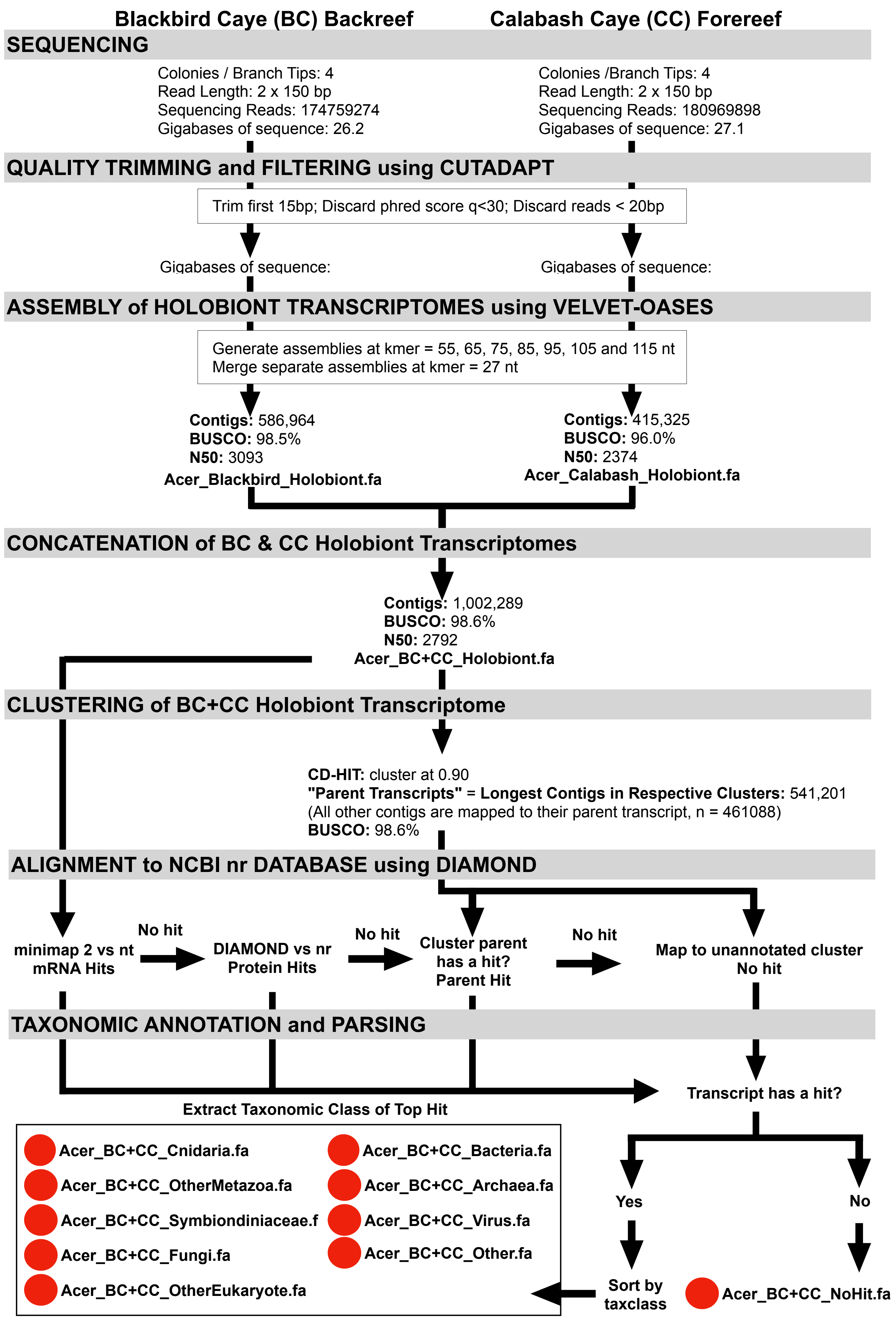

**Supplemental Figure 1.** Flow chart describing data generation and annotation. Black arrows represent steps in the data generation or analysis. Red circles denote transcript collections that are available for blast searching at https://bumpbase.bu.edu/staghorn_coral/.

**Supplemental Figure 2**. Recovery of single copy orthologs from cnidarian and Symbiodiniaceae transcriptomes.

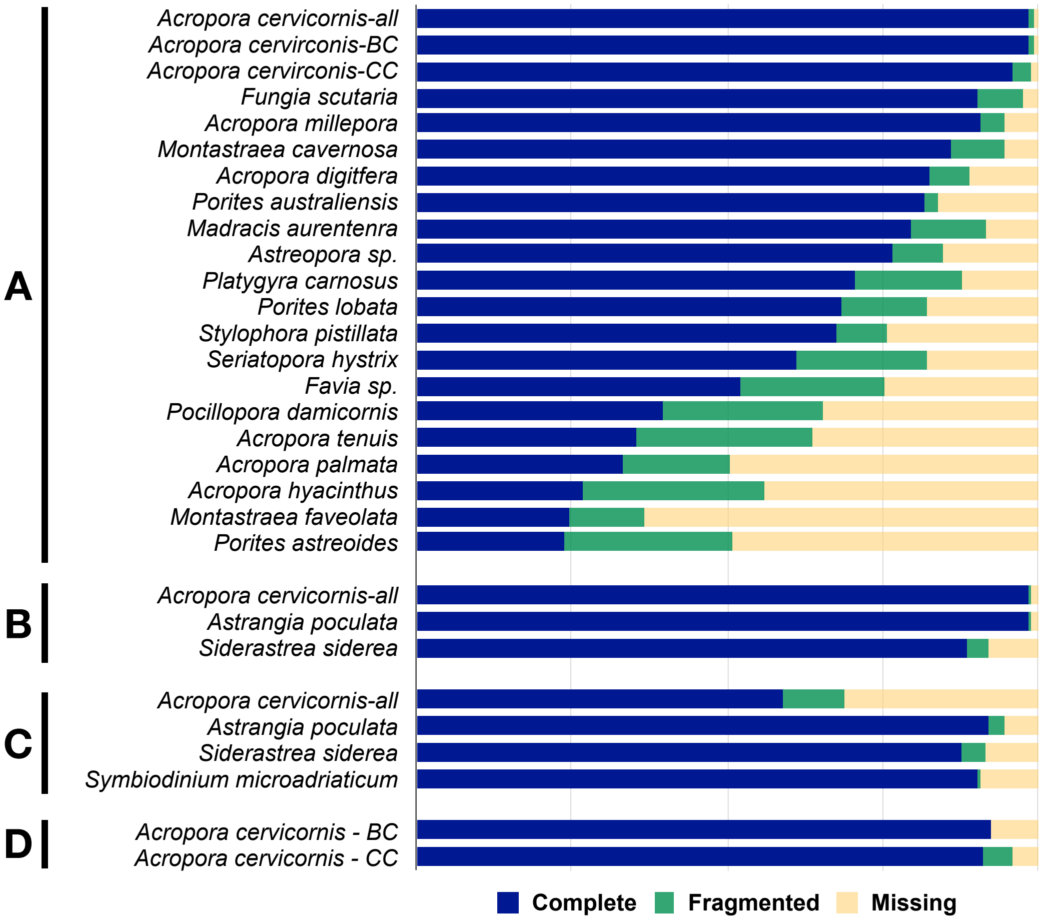

**Supplemental Figure 2.** Recovery of universal, single-copy orthologs from cnidarian and Symbiodiniaceae transcriptomes. **(A-B)** Recovery of 978 conserved metazoan orthologs from **(A)** coral holobiont transcriptomes and **(B)** coral transcript collections derived from coral-holobiont transcriptomes. **(C-D)** Recovery of 171 conserved alveolate orthologs from **(C)** coral holobiont transcriptomes and **(D)** from Symbiodiniaceae transcript collections derived from coral-holobiont transcriptomes.

**Supplemental Figure 3**. Alignment of Symbiodiniaceae actin proteins

```
Calabash MZ501686  MGDEEVAALVVDNGSGMCKAGFAGDDAPRAVFPSIVGRPKMPGIMVGMDQKDSYVGDEAQ
Blackbird MZ501688 MGDEEVAALVVDNGSGMCKAGFAGDDAPRAVFPSIVGRPKMPGIMVGMDQKDSYVGDEAQ
BAE79386.1         MGDEEVAALVVDNGSGMCKAGFAGDDAPRAVFPSIVGRPKMPGIMVGMDQKDSYVGDEAQ
Blackbird MZ501687 -MSDEVVALVVDNGSGMCKAGFSGDDAPRTVFPSLIGRPKHQSIVVGMDQRDHYVGDEAQ
BAE79388.1         -MSDEVVALVVDNGSGMCKAGFSGDDAPRTVFPSIIRRPKMPGIMVGMDQKDHLVGDEAQ
Calabash MZ501689  -MSDEVVALVVDNGSGMCKAGFSGDDAPRTVFPSIIGRPKMPGIMVGMDQKDHLVGDEAQ
                     .:**.***************:******:****:: ***  .*:*****.*  ******

Calabash MZ501686  SKRGVLTLKYPIEHGIVTNWDDMEKIWHHTFYNELRVAPEEHPVLLTEAPLNPKANRERM
Blackbird MZ501688 SKRGVLTLKYPIEHGIVTNWDDMEKIWHHTFYNELRVAPEEHPVLLTEAPLNPKANRERM
BAE79386.1         SKRGVLTLKYPIEHGIVTNWDDMEKIWHHTFYNELRVAPEEHPVLLTEAPLNPKANRERM
Blackbird MZ501687 AKRGVLALKYPIEHGIVTNWDDMERIWHHTFYNELRVSPEEHPVLLTEAPLNPKANRERM
BAE79388.1         AKRGVLALKYPIEHGIVTNWDDMERIWHHTFYNELRASPEEHPVLLTEAPLNPKANRERM
Calabash MZ501689  AKRGVLALKYPIEHGIVTNWDDMERIWHHTFYNELRVSPEEHPVLLTEAPLNPKANRERM
                   :*****:*****************.***********.:**********************

Calabash MZ501686  TQIMFETFNVPAMYVAIQAVLSLYASGRTTGIVMDSGDGVSHTVPIYEGYALPHAILRLD
Blackbird MZ501688 TQIMFETFNVPAMYVAIQAVLSLYASGRTTGIVMDSGDGVSHTVPIYEGYALPHAILRLD
BAE79386.1         TQIMFETFNVPAMYVAIQAVLSLYASGRTTGIVMDSGDGVSHTVPIYEGYALPHAILRLD
Blackbird MZ501687 TQIMFETFGVPAMYVAIQAVLSLYSSGRTTGIVTDSGDGVSHTVPIYEGYALPHAIARLD
BAE79388.1         TQIMFETFGVPAMYVAIQAVLSLYSSGRTTGIVMDSGDGVSHTVPIYEGYALPHAIARLD
Calabash MZ501689  TQIMFETFGVPAMYVAIQAVLSLYSAGRTTGIVMDSGDGVSHTVPVYEGYALPHAIARLD
                   ********.***************::******* ***********:********** ***

Calabash MZ501686  LAGRDLTEYMMKILTERGYSFTTTAEREIVRDVKEKLCYIALDFDSEMKAASESSDKEKT
Blackbird MZ501688 LAGRDLTEYMMKILTERGYSFTTTAEREIVRDVKEKLCYIALDFDSEMKAASESSDKEKT
BAE79386.1         LAGRDLTEYMVKILTERGYSFTTTAEREIVRDVKEKLCYIALDFDSEMKAASESSDKEKT
Blackbird MZ501687 LAGRDLTEYMMKIMTESGYSFSNSAEREIVRDIKEKLCYIALDFNSELKSAAESSDGAKT
BAE79388.1         LAGRDLTEYLMKIMMESGYSFSSSAERTIVRDIKEKLCYIALDFDSEMKSAAESSDKAKT
Calabash MZ501689  LAGRDLTEYLMKIMMESGYSFSSSAEREIVRDIKEKLCYIALDFDSEMKSAAESSDKAKT
                   *********::**: * ****:.:*** ****:***********:**:*:*:****  **

Calabash MZ501686  YELPDGNIITVGAERFRCPEVLFQPSFVGKEASGIHDTTFQSIMKCDVDIRKDLYSNVVL
Blackbird MZ501688 YELPDGNIITVGAERFRCPEVLFQPSFVGKEASGIHDTTFQSIMKCDVDIRKDLYSNVVL
BAE79386.1         YELPDGNIITVGAERFRCPEVLFQPSFVGKEASGIHDTTFQSIMKCDVDIRKDLYSNVVL
Blackbird MZ501687 YELPDGNIVSLHSERFRCPEVLFQPSLVGKEAMGIHDTTFQSIMRCDVDIRRDLYQNVVL
BAE79388.1         YELPDGNIVSLQSERFRCPEVLFQPSLVGKEAMGIHDTTFQSIMSCDVDIRRDLYQNVVL
Calabash MZ501689  YELPDGNIVSLQSERFRCPEVLFQPSLVGKEAMGIHDTTFRSIMSCDVDIRRDLYQNVVL
                   ********::: :*************:***** *******.*** ******.***.****

Calabash MZ501686  SGGTTMFQGIGERMTKELTALAPSTMKIKVVAPPERKYSVWIGGSILSSLSTFQQMWISK
Blackbird MZ501688 SGGTTMFQGIGERMTKELTALAPSTMKIKVVAPPERKYSVWIGGSILSSLSTFQQMWISK
BAE79386.1         SGGTTMFQGIGERMTKELTALAPSTMKIKVVAPPERKYSVWIGGSILSSLSTFQQMWISK
Blackbird MZ501687 SGGSTMLPGIGERMTKELCALAPSTVKVKVIAPPERKYSVWIGGSILSSLSTFQQMWISK
BAE79388.1         SGGSTMLPGIGERMTKELCALAPSTVKVKVIAPPERKYSVWIGGSILSSLSTFQQMWISK
Calabash MZ501689  SGGSTMLPGIGERMTKELCALAPSTVKVKVIAPPERKYSVWIGGSILSSLSTFQQMWISK
                   ***:**: ********** ******:*:**:*****************************

Calabash MZ501686  GEYDESGPTIVHRKCF
Blackbird MZ501688 GEYDESGPTIVHRKCF
BAE79386.1         GEYDESGPTTVHRKCF
Blackbird MZ501687 AEYDESGPMIVHRKCI
BAE79388.1         AEYDESGPMIVHRKCI
Calabash MZ501689  AEYDESGPMIVHRKCI
                   .*******  *****:
```

**Supplemental Figure 3.** The predicted protein sequences of two distinct Symbiodiniaceae actin proteins recovered from Blackbird and Calabash were aligned with the best matches that could be identified using blastP against the non-redundant database at NCBI. Both of the top matches (BAE79386.1, BAE79388.1) were actin proteins from unspecified *Symbiodinium* species. The sequences were aligned using the program MUSCLE (version 3.8.31; [1]) as implemented on the Phylogeny.fr web server [2]. Asterisks denote positions that are strictly conserved. Colons and periods denote conservative substitutions.

**Supplemental Figure 4**. Maximum likelihood phylogeny of Symbiodiniaceae actin

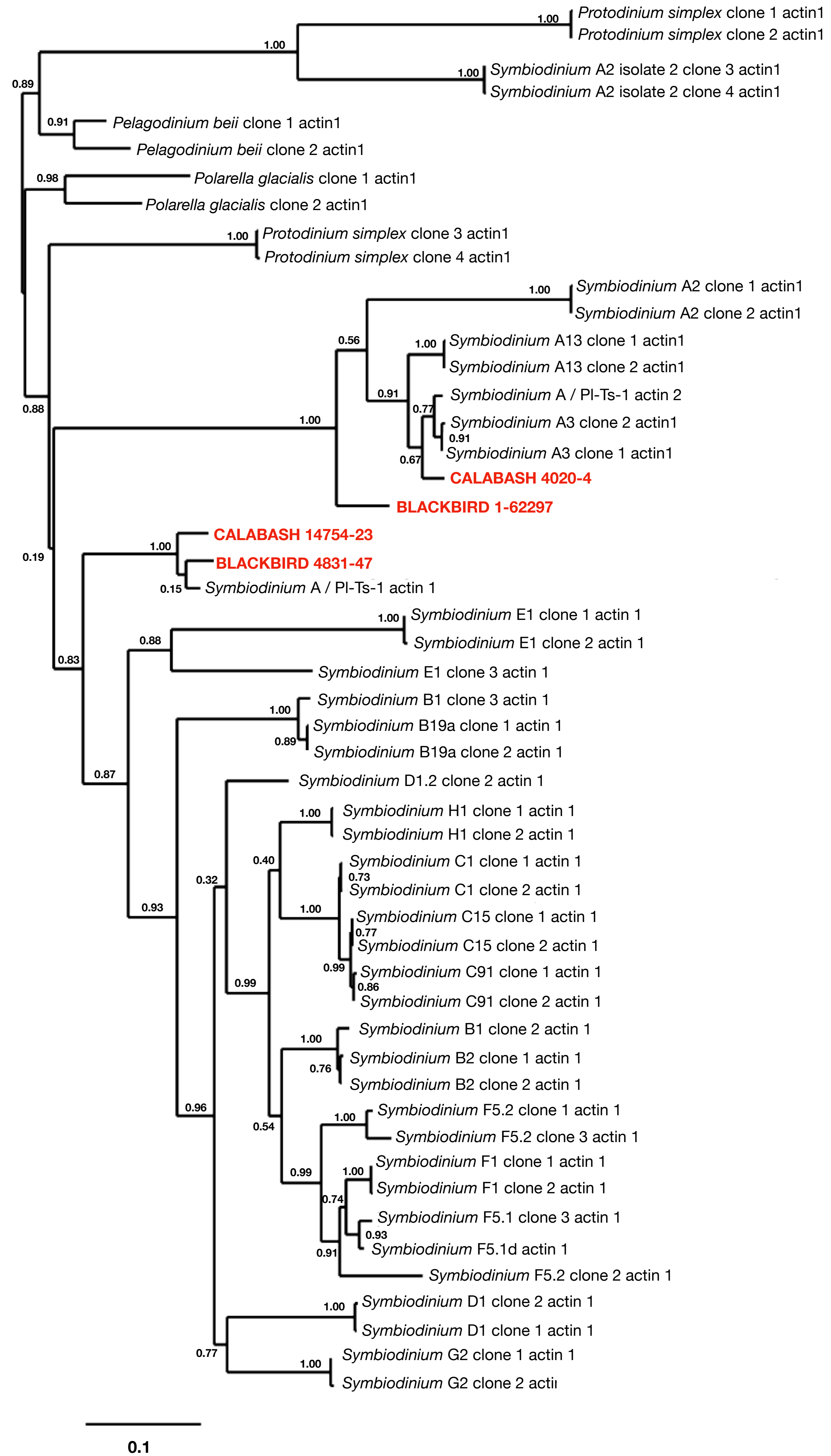

**Supplemental Figure 4.** A maximum likelihood phylogeny of Symbiodiniaceae actin nucleotide sequences. The alignment upon which the tree was based was produced using the default settings of the program MUSCLE (version 3.8.31; [1]) as implemented on the Phylogeny.fr web server [2]. Sequences from Blackbird and Calabash are shown in red (Accession numbers: MZ501686-MZ501689). The other Symbiodiniaceae actin sequences were obtained through BLAST searches conducted at NCBI or they were taken from Pochon et al. (2012) [3]. Accession numbers for these sequences are: JN558276.1, JN558277.1, JN558281.1, JN558282.1, JN558285.1, JN558286.1, JN558290.1, JN558291.1, JN558298.1, JN558299.1, JN558300.1, JN558301.1, JN558302.1, JN558303.1, JN558304.1, JN558305.1, JN558306.1, JN558307.1, JN558308.1, JN558309.1, JN558310.1, JN558313.1, JN558314.1, JN558315.1, JN558316.1, JN558317.1, JN558319.1, JN558320.1, JN558321.1, JN558322.1, JN558323.1, JN558325.1, JN558327.1, JN558329.1, JN558330.1, JN558331.1, JN558332.1, JN558334.1, JN558335.1, JN558336.1, JN558337.1, JN558338.1, JN558339.1, JN558343.1, JN558344.1, AB231899.1, AB231901.1. The alignment was then curated to retain conserved sequence blocks using Gblocks (version 0.91b [4]. The tree was generated using the default parameters of the program PhyML, v. 3.0 [5] accessed via the Phylogeny.fr web portal [2]. The tree was rendered using TreeDyn [6]. Numbers at nodes represent support for each grouping based upon approximate likelihood ratio tests [7]. The scale at lower left indicates the expected number of substitutions per nucleotide position. Parameters used in PhyML are provided below.

---

```
. Tree search : NNIs. Initial tree : BIONJ
. Model of nucleotides substitution : GTR
. Number of taxa : 51
. Log-likelihood : -9212.24145
. Discrete gamma model : Yes
  - Number of categories : 4
  - Gamma shape parameter : 0.454
. Proportion of invariant : 0.115
. Nucleotides frequencies :
  - f(A)= 0.23426
  - f(C)= 0.26780
  - f(G)= 0.27982
  - f(T)= 0.21812
. GTR relative rate parameters :

A <-> C    1.46341
A <-> G    2.81165
A <-> T    1.83521
C <-> G    1.48560
C <-> T    6.46285
G <-> T    1.00000


. Instantaneous rate matrix :

[A---------C---------G---------T------]
-0.85836   0.21305   0.42770   0.21761
 0.18636  -1.17867   0.22599   0.76633
 0.35806   0.21628  -0.69291   0.11857
 0.23371   0.94089   0.15212  -1.32671
```

**Supplemental Figure 5.** Alignment of Symbiodiniaceae COI proteins

```
Calabash MZ501690   MRIELYSSGNRIISPENQNFYNISITLHGLLMIFFLVMPGLFGGFGNYFVPIFQGSPEVV
Blackbird MZ501691  MRIELYSSGNRIISPENQNFYNVSITLHGLLMIFFLVMPGLFGGFGNYFVPIFQGSPEVV
ABK57994.1          MRIELYSSGNRIISPENQNFYNVSITLHGLLMIFFLVMPGLFGGFGNYFVPIFQGSPEVV
                    **********************:*************************************

Calabash MZ501690   YPRVNNFSILILLLSYLFLILSIISEFGGGTGWTLLPPLSTSFMTLSPSSVGNLIFGLLI
Blackbird MZ501691  YPRVNNFSILILLLSYLFLILSIISEFGGGTGWTLLPPLSTSFMTLSPSSVGNLIFGLLI
ABK57994.1          YPRVNNFSILVLLLSYLFLILSIISEFGGGTGWTLLPPLSTSFMTLSPSSVGNLIFGLLI
                    **********:*************************************************

Calabash MZ501690   SGISSCLTSLNFWITILNLRSYSLTLKTIPLFPWAFLITAFMLLLTLPVLSGTLFLVLGD
Blackbird MZ501691  SGISSCLTSLNFWVTILNLRSYSLTLKTIPLFPWAFLITALMLLLTLPVLSGTLFLVLGD
ABK57994.1          SGISSCLTSLNFWVTILNLRSYSLTLKTIPLFPWAFLITALMLLLTLPVLSGTLFLVLGD
                    *************:**************************:*******************

Calabash MZ501690   LHSNTLFFDPVFGGDPVLYQHLFWFFGHPEVYILIIPAFGVISIVISGISQLIIFGNQSM
Blackbird MZ501691  LHSNTLFFDPVFGGDPVLYQHLFWFFGHPEVYILIIPAFGVISIVISGISQLIIFGNQSM
ABK57994.1          LHSNTLFFDPVFGGDPVLYQHLFWFFGHPEVYILIIPAFGVISIVISGISQLIIFGNQSM
                    ************************************************************

Calabash MZ501690   IFAMSSISLLGSLVWGHHMYTVGLESDTRAYFTGVTILISLPTGTKIFNWLFTYLGNPPL
Blackbird MZ501691  IFAMSSISLLGSLVWGHHMYTVGLESDTRAYFTGVTILISLPTGTKIFNWLFTYLGNPPL
ABK57994.1          IFAMSSISLLGSLVWGHHMYTVGLESDTRAYFTGVTILISLPTGTKIFNWLFTYLGNPPL
                    ************************************************************

Calabash MZ501690   LHLRISSVFFSHLFLLMFTIGGSTGVILGNAAVDLALHDTYYVVAHFHFVLSLGAIISIF
Blackbird MZ501691  LHLRISSVFFSHLFLLMFTIGGSTGVILGNAAVDLGLHDTYYVVAHFHFVLSLGAIISIF
ABK57994.1          LHLRISSVFFSHLFLLMFTIGGSTGVILGNAAVDLALHDTYYVVAHFHFVLSLGAIISIF
                    ***********************************.************************

Calabash MZ501690   SGIIFNGEKIVGTKNLLFSPSSTLSLFHLHLVFVGILLTFSPMHFLGFNVMPRRIPSFPD
Blackbird MZ501691  SGIIFSGEKIVGTKSLLLSPSSTLSLFHLHLVFVGILLTFSPMHFLGFNVMPRRIPSFPD
ABK57994.1          SGIIFSGEKIVGTKNLLLSPSSTLSLFHLHLVFVGILLTFSPMHFLGFNVMPRRIPSFPD
                    *****.********.**:******************************************

Calabash MZ501690   SFNSWNSLSSIGSGITFLSFGLFYLYPFVTRSEE--------------------------
Blackbird MZ501691  SFNSWNSLSSIGSGITFLSFGLFYLYPFVTEEKARQVEWALVWWRERNEESTHLFRLSFF
ABK57994.1          SFNSWNF-----------------------------------------------------
                     ******

Calabash MZ501690   -----------
Blackbird MZ501691  HFHSIHHIILL
ABK57994.1          -----------
```

**Supplemental Figure 5.**The predicted protein sequences of Symbiodiniaceae COI proteins recovered from Blackbird and Calabash (accession numbers MZ501690, MZ501691) were aligned with the best match to both sequences that could be identified using blastP against the non-redundant database at NCBI. The top match (ABK57994.1) was a COI sequence from an unspecified *Symbiodinium* species. The sequences were aligned using the program MUSCLE (version 3.8.31; [1]) as implemented on the Phylogeny.fr web server [2]. Asterisks denote positions that are strictly conserved. Colons and periods denote conservative substitutions.

**Supplemental Figure 6.** Maximum likelihood phylogeny of Symbiodiniaceae cytochrome oxidase I

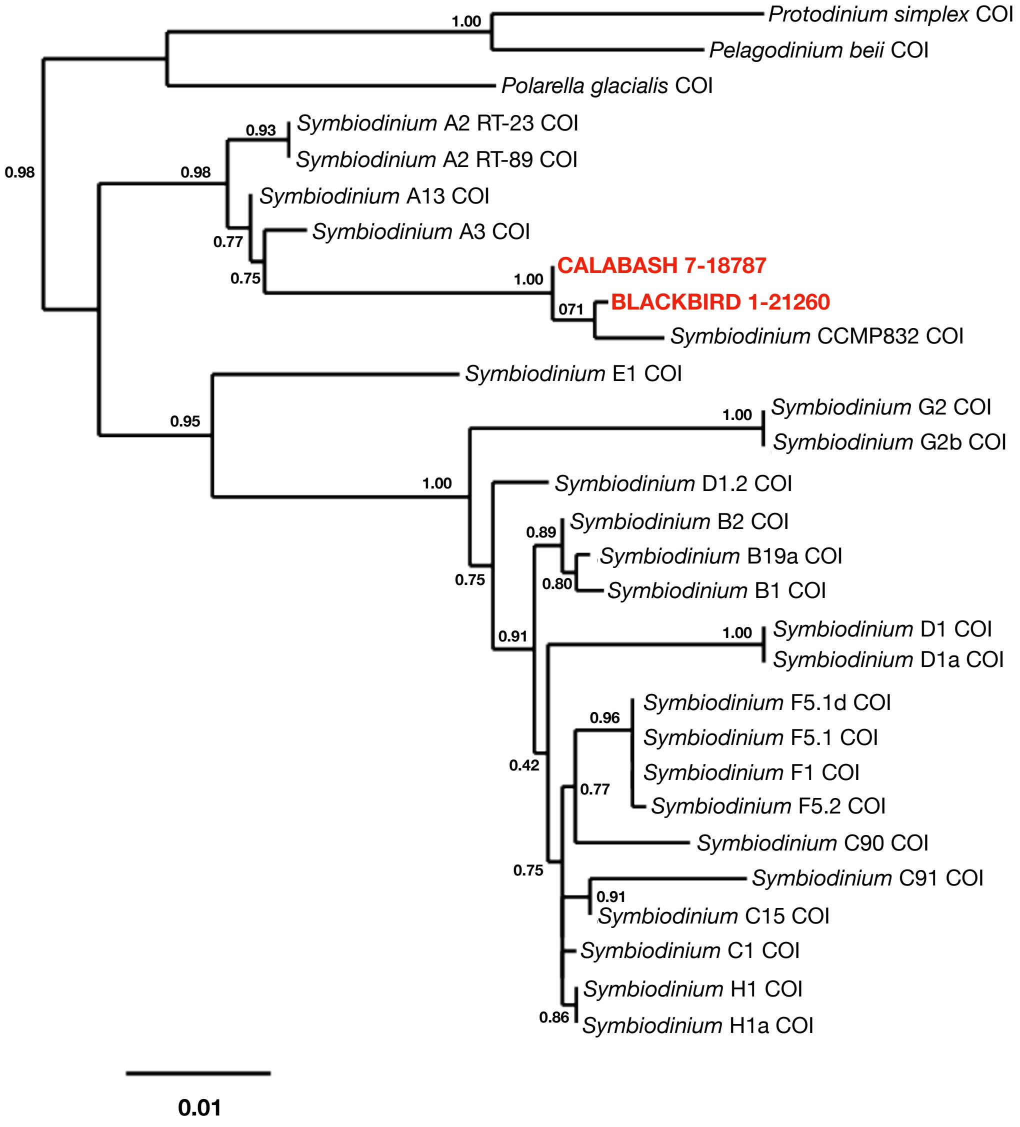

**Supplemental Figure 6.** A maximum likelihood phylogeny of Symbiodiniaceae COI nucleotide sequences. The alignment upon which the tree was based was produced using the default settings of the program MUSCLE (version 3.8.31; [1]) as implemented on the Phylogeny.fr web server [2]. Sequences from Blackbird and Calabash are shown in red (Accession numbers: MZ501690, MZ501691). The other Symbiodiniaceae actin sequences were obtained through BLAST searches conducted at NCBI or they were taken from Pochon et al. (2012) [3]. Accession numbers for these sequences are: JN557891.1, JN557892.1, JN557893.1, JN557894.1, JN557895.1, JN557896.1, JN557897.1, JN557898.1, JN557899.1, JN557900.1, JN557901.1, JN557902.1, JN557903.1, JN557904.1, JN557905.1, JN557906.1, JN557907.1, JN557908.1, JN557909.1, JN557910.1, JN557911.1, JN557912.1, JN557913.1, JN557914.1, JN557915.1, JN557916.1, EF036595.2. The alignment was then curated to retain conserved sequence blocks using Gblocks (version 0.91b [4]). The tree was generated using the default parameters of the program PhyML, v. 3.0 [5] accessed via the Phylogeny.fr web portal [2]. The tree was rendered using TreeDyn [6]. Numbers at nodes represent support for each grouping based upon approximate likelihood ratio tests [7]. The scale at lower left indicates the expected number of substitutions per nucleotide position. Parameters used in PhyML are provided below.

---

```
Tree search : NNIs. Initial tree : BIONJ
. Model of nucleotides substitution : GTR
. Number of taxa : 29
. Log-likelihood : -2958.85567
. Discrete gamma model : Yes
  - Number of categories : 4
  - Gamma shape parameter : 0.837
. Proportion of invariant : 0.362
. Nucleotides frequencies :
  - f(A)= 0.27419
  - f(C)= 0.17288
  - f(G)= 0.14094
  - f(T)= 0.41199
. GTR relative rate parameters :

A <-> C    2.27455
A <-> G    4.17649
A <-> T    1.85779
C <-> G    2.93261
C <-> T    3.33776
G <-> T    1.00000

. Instantaneous rate matrix :

[A---------C---------G---------T------]
-1.03225   0.23231   0.34776   0.45218
 0.36845  -1.42503   0.24419   0.81240
 0.67654   0.29952  -1.21946   0.24340
 0.30094   0.34091   0.08327  -0.72511
```

**Supplemental Figure 7.** Maximum likelihood phylogeny of Symbiodiniaceae ITS-2 sequences

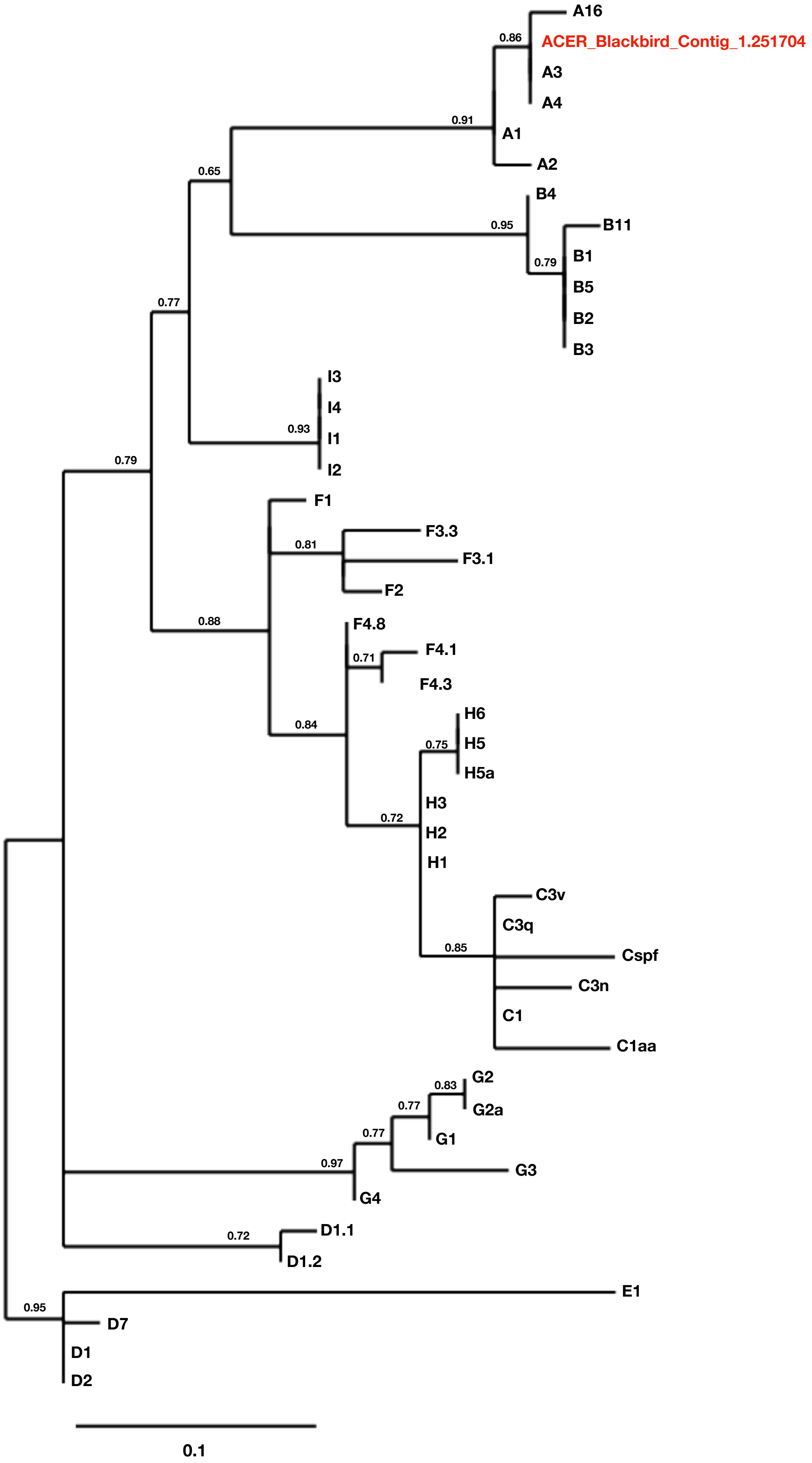

**Supplemental Figure 7.** A maximum likelihood phylogeny of Symbiodiniaceae ITS2 nucleotide sequences. The alignment upon which the tree was based was produced using the default settings of the program MUSCLE (version 3.8.31; [1]) as implemented on the Phylogeny.fr web server [2]. The ITS2 nucleotide sequences used for phylogenetic analysis include the full-length Symbiodiniaceae ITS2 sequence obtained from the Blackbird transcriptome (Accession number: MZ503278) in addition to a diversity of Symbiodiniaceae ITS2 sequences compiled by Franklin and coworkers in the GeoSymbio web application [8].

The alignment was then curated to retain conserved sequence blocks using Gblocks (version 0.91b [4]). The tree was generated using the default parameters of the program PhyML, v. 3.0 [5] accessed via the Phylogeny.fr web portal [2]. The tree was rendered using TreeDyn [6]. Numbers at nodes represent support for each grouping based upon approximate likelihood ratio tests [7]. The scale at lower left indicates the expected number of substitutions per nucleotide position. Parameters used in PhyML are provided below.

---

```
. Tree search : NNIs. Initial tree : BIONJ
. Model of nucleotides substitution : HKY85
. Number of taxa : 46
. Log-likelihood : -512.22091
. Discrete gamma model : Yes
  - Number of categories : 4
  - Gamma shape parameter : 0.592
. Proportion of invariant : 0.277
. Transition/transversion ratio : 4.799
. Nucleotides frequencies :
  - f(A)= 0.17474
  - f(C)= 0.25261
  - f(G)= 0.26445
  - f(T)= 0.30820
```

**Supplemental Figure 8.** Alignment of Symbiodiniaceae rad24 proteins

```
Blackbird MZ501693  MAAPREKSVYFAKLAEQAERYDEMAEHMKNVGSDPAELSVEERNLLSVAYKNTVGSRRAA
Blackbird MZ501692  -MPPREKDVYFAKLAEQAERYDEMADHMKNVGNADSELSVEERNLLSVAYKNTVGSRRAA
CAE7826933.1        -MPPREKDVYFAKLAEQAERYDEMADHMKNVGNADSELSVEERNLLSVAYKNTVGSRRAA
Calabash MZ501694   ---------------------------MKNVGNADSELSVEERNLLSVAYKNTVGSRRAA
                                               *****.  :************************

Blackbird MZ501693  ------------------------------------------------------------
Blackbird MZ501692  ------------------------------------------------------------
CAE7826933.1        WRIITSVMQKLGKETTKGNTENAAYAKEYCKKVEDELQSICDTILALLDGKLIAKASSGE
Calabash MZ501694   ------------------------------------------------------------


Blackbird MZ501693  ------------------------------------------------------------
Blackbird MZ501692  ------------------------------------------------------------
CAE7826933.1        SKVFYQKMKADYYRYIAEFRDGDAKSSAAEKARQAYQEAQAEDVAKKDLAVTHPIRLGLA
Calabash MZ501694   ------------------------------------------------------------


Blackbird MZ501693  ------------------------------------------------------------
Blackbird MZ501692  ------------------------------------------------------------
CAE7826933.1        LNYSVFMYEVLGNPDDACKMARTAFEDAIAELDNVAEDSYKDSTLIMQLLRDNLTLWTSD
Calabash MZ501694   ------------------------------------------------------------


Blackbird MZ501693  ------------------------------------------------------------
Blackbird MZ501692  ------------------------------------------------------------
CAE7826933.1        QEGGRVLCHSAVKQPAKLAEQAERYDEMADHMKNVGNADSELSVEERNLLSVAYKNTVGS
Calabash MZ501694   ------------------------------------------------------------


Blackbird MZ501693  ----WRIITSVMQKETTKGNSDNAAYAKEYCKKVEDELQKICDTILALLDGQLIAKASSG
Blackbird MZ501692  ----WRIITSVMQKETTKGNTENAAYAKEYCKKVEDELQSICDTILALLDGKLIAKASSG
CAE7826933.1        RRAAWRIITSVMQKETTKGNTENAAYAKEYCKKVEDELQSICDTILALLDGKLIAKASSG
Calabash MZ501694   ----WRIITSVMQKETTKGNTENAAYAKEYCKKVEDELQSICDTILALLDGKLIAKASSG
                        ****************::*****************.************:*******

Blackbird MZ501693  ESKVFYQKMKADYYRYIAEFRDGDAKSSAAEKARQAYQEAEDIAKKDLAVTHPIRLGLAL
Blackbird MZ501692  ESKVFYQKMKADYYRYIAEFRDGDAKSSAAEKARQAYQEAEDVAKKDLAVTHPIRLGLAL
CAE7826933.1        ESKVFYQKMKADYYRYIAEFRDGDAKSSAAEKARQAYQEAEDVAKKDLAVTHPIRLGLAL
Calabash MZ501694   ESKVFYQKMKADYYRYIAEFRDGDAKSSAAEKARQAYQEAEDVAKKDLAVTHPIRLGLAL
                    ******************************************:*****************

Blackbird MZ501693  NYSVFMYEVLGNPDDACKMARTAFEDAIAELDNVAEDSYKDSTLIMQLLRDNLTLWTSDQ
Blackbird MZ501692  NYSVFMYEVLGNPEDACKMARTAFEDAIAELDNVAEDSYKDSTLIMQLLRDNLTLWTSDQ
CAE7826933.1        NYSVFMYEVLGNPDDACKMARTAFEDAIAELDNVAEDSYKDSTLIMQLLRDNLTLWTSDQ
Calabash MZ501694   NYSVFMYEVLGNPEDACKMARTAFEDAIAELDNVAEDSYKDSTLIMQLLRDNLTLWTSDQ
                    *************:**********************************************

Blackbird MZ501693  EGGQA
Blackbird MZ501692  EGGQA
CAE7826933.1        EGGQA
Calabash MZ501694   EGGQA
      *****
```

**Supplemental Figure 8.** The predicted protein sequences of three distinct Symbiodiniaceae rad24 proteins recovered from Blackbird (MZ501692, MZ501693) and Calabash (MZ501694) were aligned with the best match that could be identified using blastP against the non-redundant database at NCBI. The top match (CAE7826933.1) was annotated as “artA” from an unspecified *Symbiodinium* species. ArtA, like rad24, is a member of the 14-3-3 protein family. There is an extensive region of sequence identity (indicated by the asterisks) between this top blast match and the *A. cervicornis* sequences. The sequences were aligned using the program MUSCLE (version 3.8.31; [1]) as implemented on the Phylogeny.fr web server [2]. Colons and periods denote conservative substitutions.

**Supplemental Figure 9.** Maximum likelihood phylogeny of Symbiodiniaceae rad24 proteins

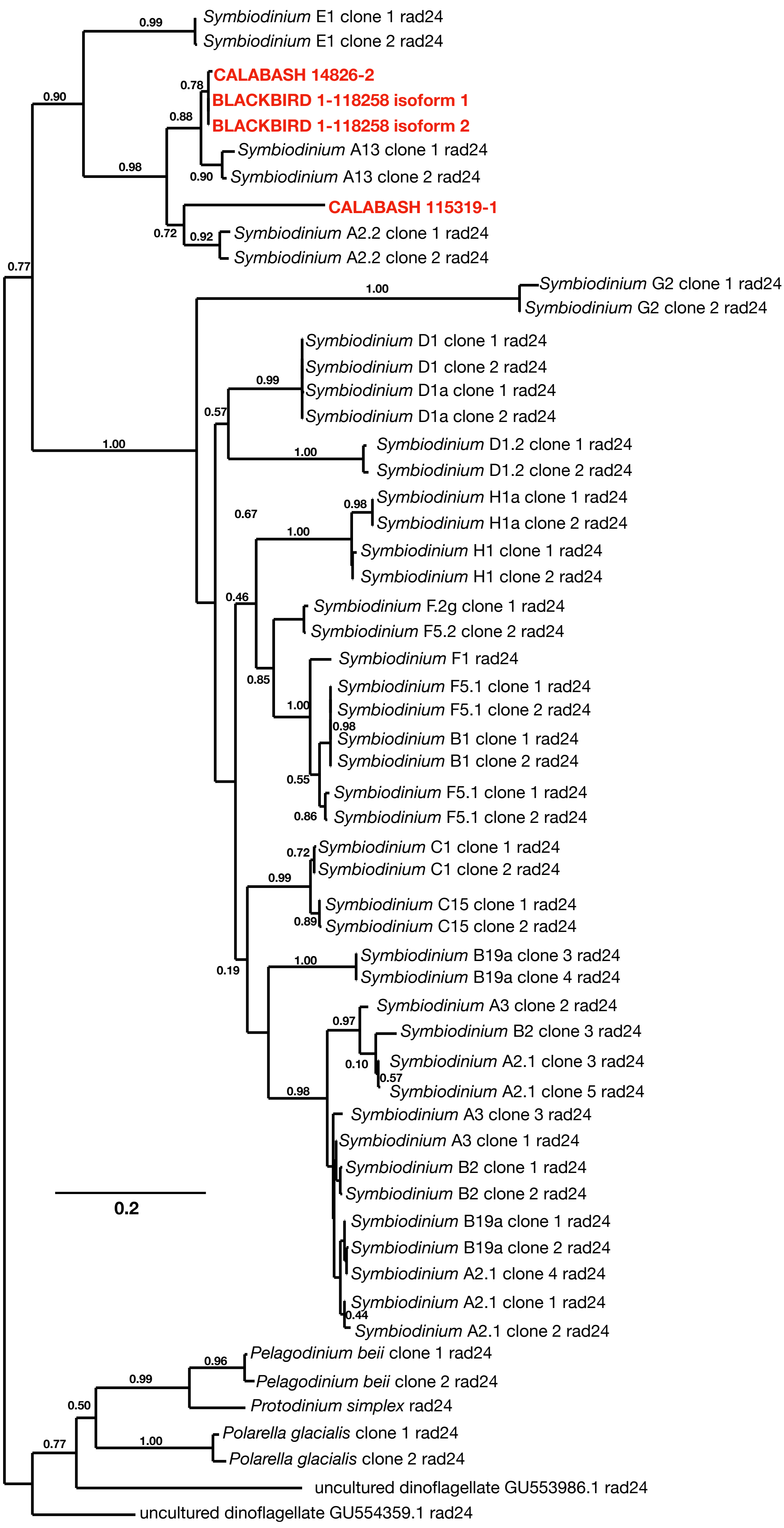

**Supplemental Figure 9.** A maximum likelihood phylogeny of Symbiodiniaceae COI nucleotide sequences. The alignment upon which the tree was based was produced using the default settings of the program MUSCLE (version 3.8.31; [1]) as implemented on the Phylogeny.fr web server [2]. Sequences from Blackbird and Calabash are shown in red (Accession numbers: MZ501690, MZ501691). The other Symbiodiniaceae actin sequences were obtained through BLAST searches conducted at NCBI or they were taken from Pochon et al. (2012) [3]. Accession numbers for these sequences are: JN558203.1, JN558204.1, JN558206.1, JN558207.1, JN558208.1, JN558209.1, JN558211.1, JN558212.1, JN558213.1, JN558214.1, JN558215.1, JN558216.1, JN558219.1, JN558220.1, JN558221.1, JN558223.1, JN558224.1, JN558229.1, JN558230.1, JN558231.1, JN558232.1, JN558233.1, JN558234.1, JN558235.1, JN558236.1, JN558237.1, JN558239.1, JN558240.1, JN558242.1, JN558243.1, JN558244.1, JN558245.1, JN558248.1, JN558249.1, JN558251.1, JN558252.1, JN558253.1, JN558254.1, JN558255.1, JN558256.1, JN558257.1, JN558259.1, JN558260.1, JN558261.1, JN558264.1, JN558265.1, JN558267.1, JN558268.1, JN558269.1, JN558272.1, JN558273.1, GU554359.1, GU553986.1. The alignment was then curated to retain conserved sequence blocks using Gblocks (version 0.91b [4]). The tree was generated using the default parameters of the program PhyML, v. 3.0 [5] accessed via the Phylogeny.fr web portal [2]. The tree was rendered using TreeDyn [6]. Numbers at nodes represent support for each grouping based upon approximate likelihood ratio tests [7]. The scale at lower left indicates the expected number of substitutions per nucleotide position. Parameters used in PhyML are provided below.

---

```
. Tree search : NNIs. Initial tree : BIONJ
. Model of nucleotides substitution : HKY85
. Number of taxa : 57
. Log-likelihood : -4709.82104
. Discrete gamma model : Yes
  - Number of categories : 4
  - Gamma shape parameter : 0.830
. Proportion of invariant : 0.312
. Transition/transversion ratio : 2.916
. Nucleotides frequencies :
  - f(A)= 0.25881
  - f(C)= 0.26258
  - f(G)= 0.27657
  - f(T)= 0.20204
```

**Supplemental Figure 10.** Alignment of coral NF-kB proteins

```
NFkB_Astrangia.               MATNSERQINETMTDSLLLDILTPGYLPDITALQMPAATYTGPYMEILEQPKQRGFRFRY
NFKB_ADIG_XP_015766406.1      MATNSEQQVNATLTDSLLMDVLNPGYLPDISALQVPTGSYQGPYLEILEQPKQRGFRFRY
NFKB_ACER_CC1_Locus_7         MATNSEQQVNATLTDSLLMDVLNPGYLPDISALQVPTGSYQGPYLEILEQPKQRGFRFRY
NFKB_ACER_BB1_Locus_1         MATNSEQQVNATLTDSLLMDVLNPGYLPDISALQVPTGSYQGPYLEILEQPKQRGFRFRY
NFKB_AMIL_XP_029186950.1      MATNSEQQVNATLTDSLLMDVLNPGYLPDISALQVPTGSYQGPYLEILEQPKQRGFRFRY
                              ******.*:* *:*****:*:*.*******:***:*:.:* ***:***************

NFkB_Astrangia                PCEGPSHGGLPGQYSEKGKKSYPSVQLCNYQGPARIVVSLVTVDEPPMPHAHSLIGKNSN
NFKB_ADIG_XP_015766406.1      PCEGPSHGGLPGEYSEKGKKSYPSVQLCNYKGPARIVVSLVTSDENPMPHAHSLIGKNAS
NFKB_ACER_CC1_Locus_7         PCEGPSHGGLPGEYSEKGKKSYPSVQLCNYKGPARIVVSLVTCDENPMPHAHSLIGKNAS
NFKB_ACER_BB1_Locus_1         PCEGPSHGGLPGEYSEKGKKSYPSVQLCNYKGPARIVVSLVTCDENPMPHAHSLIGKNAS
NFKB_AMIL_XP_029186950.1      PCEGPSHGGLPGEYSEKGKKSYPSVQLCNYKGPARIVVSLVTSDENPMPHAHSLIGKNAS
                              ************:*****************:*********** ** ************:.

NFkB_Astrangia                NGVVTVQIGPEHGMTASFPNLGIQHVTKKLVAKVLMERYVKMQSLHTATLNALNADGKAF
NFKB_ADIG_XP_015766406.1      NGVVTVQIGPEQGMTASFPNLGIQHVTKKNVGYVLMDRYLKMQMLHTATLNALTTDPRVF
NFKB_ACER_CC1_Locus_7         NGVVTVQIGPEQGMTASFPNLGIQHVTKKNVGYVLMDRYLKMQMLHTATLNALTTDPRVF
NFKB_ACER_BB1_Locus_1         NGVVTVQIGPEQGMTASFPNLGIQHVTKKNVGYVLMDRYLKMQMLHTATLNALTTDPRVF
NFKB_AMIL_XP_029186950.1      NGVVTVQIGPEQGMTASFPNLGIQHVTKKNVGLVLMDRYLKMQMLHTATLNALTIDPRVF
                              ***********:***************** *. ***:**:*** *********. * ..*

NFkB_Astrangia                DVGMLGDQALADGERATFDKNMAEAVAEEESRKVRRMVEEQTQNMNLNAVRLCFQAYLPD
NFKB_ADIG_XP_015766406.1      DIGAMVDEATADGDRGEFDKQIADTIAEEEASKVRSLVEKQKNSMNLSVVRLCFQAYLPD
NFKB_ACER_CC1_Locus_7         DIGAMVDEATADGDRGEFDKQIADTIAEEEASKVRSLVEKQKNSMNLSVVRLCFQAYLPD
NFKB_ACER_BB1_Locus_1         DIGAMVDEATADGDRGEFDKQIADTIAEEEASKVRSLVEKQKNSMNLSVVRLCFQAYLPD
NFKB_AMIL_XP_029186950.1      DIGAMVDEATADGDRGEFDKQIADTIAEEEASKVRSLVEKQKNSMNLSVVRLCFQAYLPD
                              *:* : *:* ***:*. ***::*:::****: *** :**:*.:.***..***********

NFkB_Astrangia                DSGCFTKALPPCVSNPVYDSKAPCASNLKICRMDRNSGCATGGDEVYLLCDKVQKDDIEV
NFKB_ADIG_XP_015766406.1      DNGCFTKALPPCFSRSVYDSKAPSAANLKICRMDRNSGCVTGNDEVYLLCDKVQKDYSTL
NFKB_ACER_CC1_Locus_7         DNGCFTKALPPCFSRSVYDSKAPSAANLKICRMDRNSGCVTGSDEVYLLCDKVQKDDIAV
NFKB_ACER_BB1_Locus_1         DNGCFTKALPPCFSRSVYDSKAPSAANLKICRMDRNSGCVTGSDEVYLLCDKVQKDDIAV
NFKB_AMIL_XP_029186950.1      DNGCFTKALPPCFSRSVYDSKAPSAANLKICRMDRNSGCVTGNDEVYLLCDKVQKDDIAV
                              *.**********.*..*******.*:*************.**.*************   :

NFkB_Astrangia                ---VFYETNVETAKKTWEGQGVFAPTDVHRQVAIVFKTPPYWNLAIERPAKVQLELRRKS
NFKB_ADIG_XP_015766406.1      QLCLFFNCQ-LACLTAMGGD----------TVAIVFKTPAYWNIAVERPVKVHLELRRKS
NFKB_ACER_CC1_Locus_7         ---VFYEID-ENGKRSWEGLGLFAPTDVHRQVAIVFKTPAYWNIAVERPVKVHLELRRKS
NFKB_ACER_BB1_Locus_1         ---VFYEID-ENGKRSWEGLGLFAPTDVHRQVAIVFKTPAYWNIAVERPVKVHLELRRKS
NFKB_AMIL_XP_029186950.1      ---VFYEID-ENGKRSWEGQGLFAPTDVHRQVAIVFKTPAYWNIAVERPVKVHLELRRKS
                                 :*:: :      :  *            ********.***:*:***.**:*******

NFkB_Astrangia                DQETSEPVEFTYQPQMFDKEQIGAKRRKKIPHFSDYFGSGGGGGPPPSAGGAGGGGFGGF
NFKB_ADIG_XP_015766406.1      DEETSDPVEFTYQPQMFDKEQIGSKRRKKIPHFSDYFKDGSGGGDPPAAGGGGGGGF-SF
NFKB_ACER_CC1_Locus_7         DDETSDPVEFTYQPQMFDKEQIGSKRRKKIPHFSDYFKDGSGGGDPPTAGGGGGGGF-SF
NFKB_ACER_BB1_Locus_1         DDETSDPVEFTYQPQMFDKEQIGSKRRKKIPHFSDYFKDGSGGGDPPTAGGGGGGGF-SF
NFKB_AMIL_XP_029186950.1      DDETSDPVEFTYQPQMFDKEQIGSKRRKKIPHFSDYFKDGSGGGDPPAAGGGGGGGF-SF
                              *:***:*****************:************* .*.*** **:***.***** .*

NFkB_Astrangia                GALGLLGPAAFGFIPFSTNPGTSQTNNQGGGSTSQQGQSHASGQTHTAGQSSQADLSELA
NFKB_ADIG_XP_015766406.1      PTFMFP-----GLLSFQQNPGTSQSSYQAS------------------------------
NFKB_ACER_CC1_Locus_7         STFMFP-----GLLSFQQNPSTSQSSYQGGAGTSQEGEANESGQR----QDKELSLTELA
NFKB_ACER_BB1_Locus_1         STFMFP-----GLLSFQQNPSTSQSSYQGGAGTSQEGEANESGQR----QDKELSLTELA
NFKB_AMIL_XP_029186950.1      PTFMFP-----GLLSFQQNPGTSQSSYQGGAGTSQEGEANESGQR----QDKELSLTELA
                               :: :      *::.*. **.***:. *..
```

**Supplmental Figure 10 (continued).** Alignment of coral and human NF-kB proteins

```
NFkB_Astrangia                WNLAEKSAAAMRDYAATGDMRYLLAAQRHLTAVQDDSGDTALHLAVINSQQEVVQCLIDV
NFKB_ADIG_XP_015766406.1      ------------------------------------HQNRALHLAVINSQQEVIHCLVDV
NFKB_ACER_CC1_Locus_7         WNVAAICAAAVRDYAASGDMRYLLAVQRHLTAVQDDNGDTALHLAVINSQQEVIHCLVDV
NFKB_ACER_BB1_Locus_1         WNVAAICAAAVRDYAASGDMRYLLAVQRHLTAVQDDNGDTALHLAVINSQQEVIHCLVDV
NFKB_AMIL_XP_029186950.1      WNVAAICAAAVRDYAASGDMRYLLVVQRHLTAVQDDNGDTALHLAVINSQQEVIHCLVDV
                                                                    : *************::**:**

NFkB_Astrangia                MAGLPDSYISEYNFLRQSPLHLAAITKQPRILECLLRASANARSRDRHGNTAVHIACMHG
NFKB_ADIG_XP_015766406.1      MAGLPESFVNEYNFLRQTPLHLAVITKQPRALDCLIKAGANPRLRDRHGNTAVHIACSYG
NFKB_ACER_CC1_Locus_7         MAGLPESFVNEYNFLRQTPLHLAVITKQPRALDCLIKAGANPRLRDRHGNTAVHIACSYG
NFKB_ACER_BB1_Locus_1         MAGLPESFVNEYNFLRQTPLHLAVITKQPRALDCLIKAGANPRLRDRHGNTAVHIACSYG
NFKB_AMIL_XP_029186950.1      MAGLPESFVNEYNFLRQTPLHLAVITKQPRALDCLIKAGANPRLRDRHGNTAVHIACSYG
                              *****:*::.*******:*****.****** *:**:.*.**.* ************* :*

NFkB_Astrangia                DAVCLKALLNFNVSKTVLNWQNYQGLTPVHLAVLAGSKDVLKLLNSAGANMSAQDGTSGK
NFKB_ADIG_XP_015766406.1      DATCLKALLHYDVSKMVLNWQNYQGLTPVHLAVLCGSKDVLKLLRSAGANMSAQDGTSGK
NFKB_ACER_CC1_Locus_7         DATSLKALLHYDVSKMVLNWQNYQGLTPVHLAVLCGSKDVLKLLRSAGANMSSQDGTSGK
NFKB_ACER_BB1_Locus_1         DATSLKALLHYDVSKMVLNWQNYQGLTPVHLAVLCGSKDVLKLLRSAGANMSSQDGTSGK
NFKB_AMIL_XP_029186950.1      DATCLKALLHYDVSKMVLNWQNYQGLTPVHLAVLCGSKDVLKLLRSAGANMSAQDGTSGK
                              **..*****:::*** ******************.*********.*******:*******

NFkB_Astrangia                TPLHHAVEQDHLAVAGFLILEANSDVDAATFDGNTPLHIAAASGLKGQTALLVAAGADTT
NFKB_ADIG_XP_015766406.1      TPLHLSVEQDNLSLSGFLILEANCDVDASTFDGNTPLHLAAGLGLKGHTALLVAAGADTT
NFKB_ACER_CC1_Locus_7         TPLHLSVEQDNLSLSGFLILEANCDVDASTFDGNTPLHLAAGLGLKGHTALLVAAGADTT
NFKB_ACER_BB1_Locus_1         TPLHLSVEQDNLSLSGFLILEANCDVDASTFDGNTPLHLAAGLGLKGHTALLVAAGADTT
NFKB_AMIL_XP_029186950.1      TPLHLSVEQDNLSLSGFLILEANCDVDASTFDGNTPLHLAAGLGLKGHTALLVAAGADTT
                              **** :****:*:::********.****:*********:**. ****:************

NFkB_Astrangia                LQNSDEETAFDLANVAEVQEILDEDEALSTDPSQDDELTAGLTGLTLGQGDMEKLDPYVR
NFKB_ADIG_XP_015766406.1      FPNSEDETAFDLANVAEVQEILNEDEAPSGDPV--EELETGVISIRLGKGDLDQLDPYVR
NFKB_ACER_CC1_Locus_7         FPNSEDETAFDLANVAEVQEILDEDEAPSADPV--EELETGVISIRLGKGDLDQLDPYVR
NFKB_ACER_BB1_Locus_1         FPNSEDETAFDLANVAEVQEILDEDEAPSADPV--EELETGVISIRLGKGDLDQLDPYVR
NFKB_AMIL_XP_029186950.1      FPNSEDETAFDLANVAEVQEILDEDEAPSADPV--EELETGVISIRLGKGDLDQLDPYVR
                              : **::****************:**** * **   :** :*: .: **:**:::******

NFkB_Astrangia                RKMAQRLDPSSSTGADWRELARRLGLGTLETAFAIHSSPTTQVLAQYEAADGSIKTLRQV
NFKB_ADIG_XP_015766406.1      RKMAQRLDP--STGADWRDFARRLGLGTLANAFAIQSSPTVQVLVHFEAADGTMEKLREV
NFKB_ACER_CC1_Locus_7         RKMAQRLDP--STGADWRDLARRLGLGTLANAFAIQSSPTVQVLAHFEAADGTMEKLREV
NFKB_ACER_BB1_Locus_1         RKMAQRLDP--STGADWRDLARRLGLGTLANAFAIQSSPTVQVLAHFEAADGTMEKLREV
NFKB_AMIL_XP_029186950.1      RRMAQRLDP--STGADWRDLARRLGLGTLANAFAIQSSPTVQVLAHFEAADGTMEKLREV
                              *.*******  *******::********* .****:****.***.::*****:::.**:*

NFkB_Astrangia                LHDMRRGDVLEILEE--RNDSGFDSGLGSHSLSASRSIEMASYEAGSSNISSSSSIQKGT
NFKB_ADIG_XP_015766406.1      LYDMRRGDVLEILDQESRHDSGFDSGFGSQSLSINRSEGDTSSGAHSSDVASRSSLSKDF
NFKB_ACER_CC1_Locus_7         LYDMRRGDVLEILDQESRHDSGFDSGFGSQSLSINRSEGGTSSGAHSSDVASRSSLSKDF
NFKB_ACER_BB1_Locus_1         LYDMRRGDVLEILDQESRHDSGFDSGFGSQSLSINRSEGGTSSGAHSSDVASRSSLSKDF
NFKB_AMIL_XP_029186950.1      LYDMRRGDVLEILDQESRHDSGFDSGFGSQSLSINRSEGDTSSGAHSSDVASRSSLSKDF
                              *:***********::  *:*******:**:*** .**   :*  * **:::* **:.*.

NFkB_Astrangia                TSLESTRKAGNFQPIRQHQDVF
NFKB_ADIG_XP_015766406.1      HKLESSQESAKTRPLLQHQHVF
NFKB_ACER_CC1_Locus_7         HKLESSQESAKTRPLLQQQHVF
NFKB_ACER_BB1_Locus_1         HKLESSQESAKTRPLLQQQHVF
NFKB_AMIL_XP_029186950.1      HKLESSQESAKTRPLLQQQHVF
                               .***:.::.: .*: *:* **
```

**Supplemental Figure 10.** NF-kB protein sequences from *A. cervicornis* (ACER; Blackbird & Calabash) aligned to NF-kB sequences from other *Acropora* species (ADIG=*A. digitifera*; AMIL=*A. millepora*) and the temperate coral *Astrangia poculata* [9]. The Blackbird and Calabash sequences are identical to each other and identical to the *A. millepora* sequence at 891of 904 residues. The sequences were aligned using the program MUSCLE (version 3.8.31; [1]) as implemented on the Phylogeny.fr web server [2]. Asterisks denote positions that are strictly conserved. Colons and periods denote conservative substitutions.

**Supplemental Figure 11.** Conserved motifs in coral and human NF-kB proteins

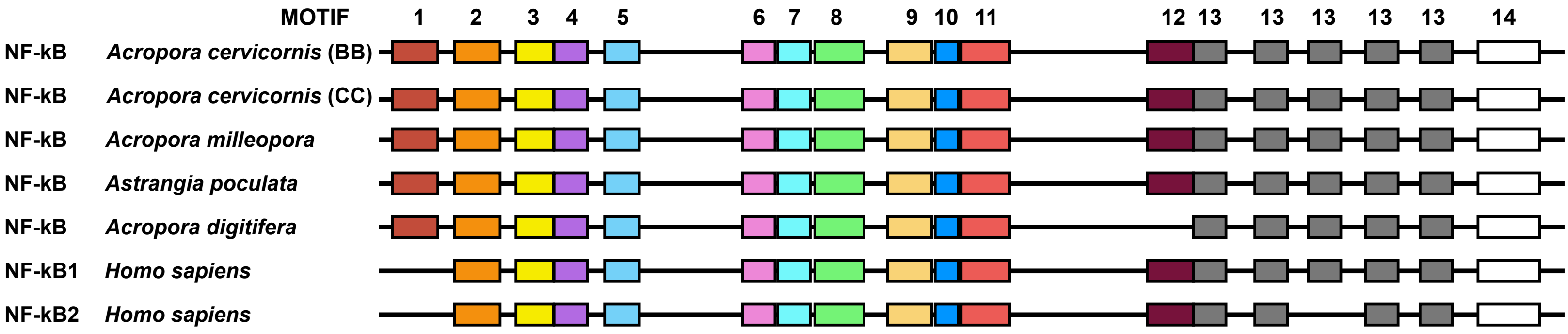


**Supplemental Figure 11.** Diagrammatic representation of NFkB proteins in corals and humans showing the location of conserved motifs. The motifs were originally identified by Nguyen in a phylogenetically diverse collection of metazoan NFkB proteins using MEME [9,10]. These previously defined motifs were located in *Acropora* NFkB proteins in the current study using MAST [11]. Motifs are numbered sequentially from the amino terminus to the carboxy terminus, as in Nguyen [9]. By definition, the length of a given motif, represented by the width of each colored box, is conserved in all instances of that motif. Intermotif distances (represented by the thin black line) are not drawn to scale.

**Supplemental Figure 12.** Alignment of coral and human IKKa/b proteins

```
IKKab APOC      MECGRERKRSREG-WVEERCLGSGAFGTVLLYENKITKEQIALKRCRMELNPHNKKLWHQ
IKKab_AMIL      MEYGRAINLSRDG-WMEERCLGAGGFGTVMLYENKITKEQVALKRCRLELNSHNRKRWQQ
IKKab_ACER_BC   MEYGRAINLSRDG-WMEEGCLGAGGFGTVMLYENKITKEQVALKRCRLELNSHNRKRWQQ
IKKab_ACER_CC   MEYGRAINLSRDG-WMEERCLGAGGFGTVMLYENKITKEQVALKRCRLELNSHNRKRWQQ
IKKA_HUMAN      MERPPGLRPGAGGPWEMRERLGTGGFGNVCLYQHRELDLKIAIKSCRLELSTKNRERWCH
IKKB_HUMAN      MSWSPSLTTQTCGAWEMKERLGTGGFGNVIRWHNQETGEQIAIKQCRQELSPRNRERWCL
                *.          * *     **:*.**.*  : :.    ::*:* ** **...*.: *

IKKab APOC      EVDIMKRLEHPNLVSAKDVPPPL-DVSDDELPLLAMEFCSGGDLRKVLNAPESCCGLKEK
IKKab_AMIL      EVEIMKRLHHPNLVSARDVPPAL-DVSDDELPLLAMEFCSGGDLRKVLNAPESCCGLKET
IKKab_ACER_BC   EVEIMKRLHHPNLVSARDVPPAL-DVSDDELPLLAMEFCSGGDLRKVLNAPESCCGLKET
IKKab_ACER_CC   EVEIMKRLHHPNLVSARDVPPAL-DVSDDELPLLAMEFCSGGDLRKVLNAPESCCGLKET
IKKA_HUMAN      EIQIMKKLNHANVVKACDVPEEL-NILIHDVPLLAMEYCSGGDLRKLLNKPENCCGLKES
IKKB_HUMAN      EIQIMRRLTHPNVVAARDVPEGMQNLAPNDLPLLAMEYCQGGDLRKYLNQFENCCGLREG
                *::**..* *.*:* * ***  : ::   ::******:*.****** **  *.****.*

IKKab APOC      TVLKITSDVAAAVEFLHGHRIIHRDLKPENIVISHADNKVVYKLIDLGYAKELDQGSLAT
IKKab_AMIL      TVLQIAGDVAAAVEFLHSHRIIHRDLKPENIVINRVDRKVIYKLIDLGYAKELDQASMAT
IKKab_ACER_BC   TVLQIAGDVAAAVEFLHSHRIIHRDLKPENIVINHVDRKVIYKLIDLGYAKELDQASMAT
IKKab_ACER_CC   TVLQIAGDVAAAVEFLHSHRIIHRDLKPENIVINHVDRKVIYKLIDLGYAKELDQASMAT
IKKA_HUMAN      QILSLLSDIGSGIRYLHENKIIHRDLKPENIVLQDVGGKIIHKIIDLGYAKDVDQGSLCT
IKKB_HUMAN      AILTLLSDIASALRYLHENRIIHRDLKPENIVLQQGEQRLIHKIIDLGYAKELDQGSLCT
                 :* : .*:.:.: :** :.************:.    .:::*:*******::**.*:.*

IKKab APOC      TFVGTFKYLAPELLDKVKYTKTVDYWTFGTVLFECITGMRPFLTELSPVQWYNKVRDKGP
IKKab_AMIL      TFVGTLKYLAPELLDRVEYTKSVDYWSFGTVLFECITGVRPFLPELSPVKWHNVVFKKGA
IKKab_ACER_BC   TFVGTLKYLAPELLDRVEYTKSVDYWSFGTVLFECITGVRPFLPELSPVKWHNVVFKKGA
IKKab_ACER_CC   TFVGTLKYLAPELLDRVEYTKSVDYWSFGTVLFECITGVRPFLPELSPVKWHNVVFKKGA
IKKA_HUMAN      SFVGTLQYLAPELFENKPYTATVDYWSFGTMVFECIAGYRPFLHHLQPFTWHEKIKKKDP
IKKB_HUMAN      SFVGTLQYLAPELLEQQKYTVTVDYWSFGTLAFECITGFRPFLPNWQPVQWHSKVRQKSE
                :****::******::.  ** :****:***: ****:* ****   .*. *:. : .*.

IKKab APOC      DDICAYFDLKEDIRFSSVLPTPNSLCRQLQDKFVALLRLLLLWDPKARGGSI-LQNGKRE
IKKab_AMIL      DDICAFFDVKGEVQFSSILPVPNTLCRQLQDKFVSLLRLLLLWDPKRRGGLV-LDNGKRE
IKKab_ACER_BC   DDICAFFDVKGEVQFSSVLPVPNTLCRQLQDKFVSLLRLLLLWDPKRRGGLV-LDNGKRE
IKKab_ACER_CC   DDICAFFDVKGEVQFSSVLPVPNTLCRQLQDKFVSLLRLLLLWDPKRRGGLV-LDNGKRE
IKKA_HUMAN      KCIFACEEMSGEVRFSSHLPQPNSLCSLVVEPMENWLQLMLNWDPQQRGGPVDLTLKQPR
IKKB_HUMAN      VDIVVSEDLNGTVKFSSSLPYPNNLNSVLAERLEKWLQLMLMWHPRQRGTD--PTYGPNG
                  * .  ::.  :.*** ** **.*   : : :   *.*:* * *. **

IKKab APOC      CFDVLEKIINTVIVRVFTISSAVVLPFEVLPTETIEDLQVKICEVTGVDVREQELLTASG
IKKab_AMIL      CFDMLEKIVNTVVVHVFPIATATLISFEVFPSETVDDLKVKLSEATGVK-EEHELLTVSG
IKKab_ACER_BC   CFDMLEKIVNTVVVHVFPIATATLISFEVFPSETVDDLKVKLSEATGVK-EEHELLTVSG
IKKab_ACER_CC   CFDMLEKIVNTVVVHVFPIATATLISFEVFPSETVDDLKVKLSEATGVK-EEHELLTVSG
IKKA_HUMAN      CFVLMDHILNLKIVHILNMTSAKIISFLLPPDESLHSLQSRIERETGINTGSQELLSETG
IKKB_HUMAN      CFKALDDILNLKLVHILNMVTGTIHTYPVTEDESLQSLKARIQQDTGIPEEDQELLQEAG
                **  :: *:*  :*.:: : :. : .: :   *:: .*: .:   **:   .:***  :*

IKKab APOC      QAVKPKTVVVD-CIKDKKTGE----DC-VLFLLPTE----NEPTSTRPFYTMPPTVQGMV
IKKab_AMIL      QLLESNGLVMDICNRDSKFEG----DC-LLFLISTG----NGSLSPPPIFALPGTVHGMV
IKKab_ACER_BC   QLLESNGLVMDICNRDSKFEG----DC-LLFLISTG----NGSLSPPPIFALPGTVHGMV
IKKab_ACER_CC   QLLESNGLVMDICNRDSKFEG----DC-LLFLISTG----NGSLSPPPIFALPGTVHGMV
IKKA_HUMAN      ISLDPRKPASQ-CVLDGVRGC----DSYMVYLFDKSKTVYEGPFASRS---LSDCVNYIV
IKKB_HUMAN      LALIPDKPATQ-CISDGKLNEGHTLDMDLVFLFDNSKITYETQISPRP---QPESVSCIL
                  : .   . : *  *         *  :::*: .     :   :. .    .  *  ::
```

**Supplemental Figure 12.** Alignment of coral and human IKKa/b proteins (continued)

```
IKKab_APOC      TEQKTLMSYEELKRALAQGVNFCREQIKLNQLLIEGFRAAQISLLVLNSTLGQLKTDLMT
IKKab_AMIL      TEPKTLMSYEDLKKAFAVGVNFCWEQRRINLLLIEGFRAAQISLLHLNSRLGQVKNDVTA
IKKab_ACER_BC   TEPKTLMSYEDLKKAFAVGVNFCWEQRRINILLIEGFRAAQISLLHLNSRLGQVKNDVTT
IKKab_ACER_CC   TEPKTLMSYEDLKKAFAVGVNFCWEQRRINILLIEGFRAAQISLLHLNSRLGQVKNDVTT
IKKA_HUMAN      QDSKIQLPIIQLRKVWAEAVHYVSGLKEDYSRLFQGQRAAMLSLLRYNANLTKMKNTLIS
IKKB_HUMAN      QEPKRNLAFFQLRKVWGQVWHSIQTLKEDCNRLQQGQRAAMMNLLRNNSCLSKMKNSMAS
                 : *  :.  :*... .   :           * :* *** :.**  *: * ::*. : :

IKKab_APOC      EFNKLEAKKDFFTSSMQTDLEIYADNINI-VKNDKLLRSWKRAQKKVESF-HNNDVPQLK
IKKab_AMIL      ELKKLEAKVDFFNSSLQFDLECYEENISL-MKNDKVFRAWKRAQKKVASF-QNNDVLALL
IKKab_ACER_BC   ELKKLEAKVDFFNSSLQFDLECYEENISL-MKNDKVFRAWKRAQKKVASF-QNNDVLALL
IKKab_ACER_CC   ELKKLEAKVDFFNSSLQFDLECYEENISL-MKNDKVFRAWKRAQKKVASF-QNNDVLALL
IKKA_HUMAN      ASQQLKAKLEFFHKSIQLDLERYSEQMTYGISSEKMLKAWKEMEEKAIHYAEVGVIGYLE
IKKB_HUMAN      MSQQLKAKLDFFKTSIQIDLEKYSEQTEFGITSDKLLLAWREMEQAVELCGRENEVKLLV
                  ::*:** :** .*:* *** * ::    :..:*:: :*.  :: .      . :  *

IKKab_APOC      DLTTALQGRVVELQKSPYASGTQSKSKMDEMRYQCAVQLYEKLKQSDRRP---ADCQQMA
IKKab_AMIL      NSTMILQGKIVESQKGPYASGIQAKSKVDGM-YNNVLKLFEDLRQGDRRP---ADSQKIV
IKKab_ACER_BC   NSTMVLQGKIVESQKGPYASGTQAKSKVDGM-YSNVLQLFEDLRQGDRRP---ADSQKIV
IKKab_ACER_CC   NSTMVLQGKIVESQKGPYASGTQAKSKVDGM-YSNVLQLFEDLRQGDRRP---ADSQKIV
IKKA_HUMAN      DQIMSLHAEIMELQKSPY--GRRQGDLMESL-EQRAIDLYKQLKHRPSDH-SYSDSTEMV
IKKB_HUMAN      ERMMALQTDIVDLQRSPM--GRKQGGTLDDL-EEQARELYRRLREKPRDQRTEGDSQEMV
                :    *:  ::: *..*   * .  . :: :  . . .*:  *.         .*. ::.

IKKab_APOC      TIILQILAQREKLHKDIFTQLSEVMTCRREMRRVMHRLLQVKAKLVSRAEELSEMQKQRQ
IKKab_AMIL      AVILQFLAQKEKLHKDILSCLSKIISCKSEVREMMIKLDKAKSKLLERQQELVEIQNQRQ
IKKab_ACER_BC   AVILQFLAQKEKLHKDILSRLSKIISCKSEIREMMIKLDKAKSKLLERQQELEEMQNQRQ
IKKab_ACER_CC   AVILQFLAQKEKLHKDILSRLSKIISCKSEIREMMIKLDKAKSKLLERQQELEEMQNQRQ
IKKA_HUMAN      KIIVHTVQSQDRVLKELFGHLSKLLGCKQKIIDLLPKVEVALSNIKEADNTVMFMQGKRQ
IKKB_HUMAN      RLLLQAIQSFEKKVRVIYTQLSKTVVCKQKALELLPKVEEVVSLMNEDEKTVVRLQEKRQ
                 :::: : . :.  . :   **: : *. :   :: .:  . : : .  : :  :* :**

IKKab_APOC      VDLWRLLQ--------------SKKQKEQDRHCSSSSSHASLSSLATSCSSSS----SHA
IKKab_AMIL      ADLWKILQ--------------SMRGQDSRRISS--GSCTSLSSPDISCT-------ESI
IKKab_ACER_BC   ADLWKILQ--------------SMRSQDSRRISS--GSCTSLSSPDISCT-------ESI
IKKab_ACER_CC   ADLWKILQ--------------SMRSQDSRRISS--GSCTSLSSPDISCT-------ESI
IKKA_HUMAN      KEIWHLLKIACTQSSARSLVGSSLEGAVTPQTSAWLPPTSAEHDHSLSCVVTPQDGETSA
IKKB_HUMAN      KELWNLLKIACSK--VRGPVSGSPDSMNASRLS---QPGQLMSQPSTASNSLPEPAKKSE
                 ::*.:*:              *       . .    .     .   :.

IKKab_APOC      SLSSLATSCIRTTDCVLEQEVASEESDACE----LLDEQ--------------
IKKab_AMIL      VLIEESSRGFEKFAEVVENLKLDQINETLNLDWEFLEEDSGPDDKKWEGRPAS
IKKab_ACER_BC   VLIEESSRGFEKFAEVVENLKLDQINETLNLDWEFLEEDSGPDDKKWEGRPAS
IKKab_ACER_CC   VLIEESSRGFEKFAEVVENLKLDQINETLNLDWEFLEEDSGPDDKKWEGRPAS
IKKA_HUMAN      QMIEENLNCLGHLSTIIHEANEEQGNSMMNLDWSWLTE---------------
IKKB_HUMAN      ELVAEAHNLCTLLENAIQDTVREQDQSFTALDWSWLQTE---EEEHSCLEQAS
                 :              : :   .: ..        *
```

**Supplemental Figure 12.** IKK a/b protein sequences from *A. cervicornis* (ACER; Blackbird & Calabash) aligned to NF-kB sequences from human, other *Acropora* species (ADIG=*A. digitifera*; AMIL=*A. millepora*), and the temperate coral *Astrangia poculata* (APOC) [9]. The Blackbird (NCBI accession MZ812722) and Calabash (MZ812723) sequences are identical to each other at 730/731 residues and identical to the *A. millepora* sequence at 717-718 of 731 residues. The sequences were aligned using the program MUSCLE (version 3.8.31; [1]) as implemented on the Phylogeny.fr web server [2]. Asterisks denote positions that are strictly conserved. Colons and periods denote conservative substitutions.

**Supplemental Figure 13.** Alignment of coral and human IKKE/TBK1 proteins

```
IKKeTBK1_APOC     MEIRSSANYIWNIKDILGQGATGAVYKGRNKKTGDTFAVKISNQLGMMRPVEVRRREFEV
IKKeTBK1_AMIL_v3  METRGSLNYQWNIRDQLGQGATGAVFKGRHKKTGETYAIKVSNNFGMMRPVEIRRREYDV
IKKeTBK1_AMIL_v6  METRGSLNYQWNIRDQLGQGATGAVFKGRHKKTGETYAIKVSNNFGMMRPVEIRRREYDV
IKKeTBK1_ACER_BC  METRGSVNYQWNIRDQLGQGATGAVFKGRHKKTGETYAIKVSNHFGMMRPVEIRRREYDV
IKKeTBK1_ACER_CC  METRGSVNYQWNIRDQLGQGATGAVFKGRHKKTGETYAIKVSNHFGMMRPVEIRRREYDV
IKKe_HUMAN        --MQSTANYLWHTDDLLGQGATASVYKARNKKSGELVAVKVFNTTSYLRPREVQVREFEV
TBK1_HUMAN        --MQSTSNHLWLLSDILGQGATANVFRGRHKKTGDLFAIKVFNNISFLRPVDVQMREFEV
                     ..: *: *   * ******. *:..*:**:*:  *:*: *  . :** ::. **::*

IKKeTBK1_APOC     LNKLNHENIVKLYASETELRSQNEIIVMELCSGGSLFTLLENPANAYGFLEDDFKQVVKD
IKKeTBK1_AMIL_v3  LLKLHHENIVKIFATEVEIRGQNEIIVMELCSGGSLFTMLENPSNAYGLQEDEFKQVVKD
IKKeTBK1_AMIL_v6  LLKLHHENIVKIFATEVEIRGQNEIIVMELCSGGSLFTMLENPSNAYGLQEDEFKQVVKD
IKKeTBK1_ACER_BC  LLKLNHENIVKIFATEVEIRGQNEIIVMELCSGGSLFTMLENPSNAYGLQEDEFKQVVKD
IKKeTBK1_ACER_CC  LLKLNHENIVKIFATEVEIRGQNEIIVMELCSGGSLFTMLENPSNAYGLQEDEFKQVVKD
IKKe_HUMAN        LRKLNHQNIVKLFAVEETGGSRQKVLVMEYCSSGSLLSVLESPENAFGLPEDEFLVVLRC
TBK1_HUMAN        LKKLNHKNIVKLFAIEEETTTRHKVLIMEFCPCGSLYTVLEEPSNAYGLPESEFLIVLRD
                  * **:*:****::* *     .:::::** *. *** ::**.* **:*: *.:*  *:.

IKKeTBK1_APOC     VAAGMKHLREQGMVHRDLKPGNIMRVAKVDGSFVYKLTDFGAARELEAHEGFMSVYGTEE
IKKeTBK1_AMIL_v3  VVAGMKHLREQGIVHRDLKPGNIMRVAKDDGSFVYKLTDFGAAKELEGSEEFMSIYGTEE
IKKeTBK1_AMIL_v6  VVAGMKHLREQGIVHRDLKPGNIMRVAKDDGSFVYKLTDFGAAKELEGSEEFMSIYGTEE
IKKeTBK1_ACER_BC  VVAGMKHLREQGIVHRDLKPGNIMRVAKDDGSFVYKLTDFGAAKELEGSEEFMSIYGTEE
IKKeTBK1_ACER_CC  VVAGMKHLREQGIVHRDLKPGNIMRVAKDDGSFVYKLTDFGAAKELEGSEEFMSIYGTEE
IKKe_HUMAN        VVAGMNHLRENGIVHRDIKPGNIMRLVGEEGQSIYKLTDFGAARELDDDEKFVSVYGTEE
TBK1_HUMAN        VVGGMNHLRENGIVHRDIKPGNIMRVIGEDGQSVYKLTDFGAARELEDDEQFVSLYGTEE
                  *..**:****:*:****:*******:   :*. :*********.**:  * *:*:*****

IKKeTBK1_APOC     YLHPDLYERGVLRKHGNKAFGAMVDLWSIGVTFYHIATGQLPFRPYGG-RQNRDTMHLIT
IKKeTBK1_AMIL_v3  YLHPDLYERGVLRKHGNKAFSATVDLWSIGVTFYHIATGQLPFRPYGG-RQNKVTMYEIT
IKKeTBK1_AMIL_v6  YLHPDLYERGVLRKHGNKAFSATVDLWSIGVTFYHIATGQLPFRPYGG-RQNKVTMYEIT
IKKeTBK1_ACER_BC  YLHPDLYERGVLRKHGNKAFSATVDLWSIGVTFYHIATGQLPFRPYGG-RQNKVTMYEIT
IKKeTBK1_ACER_CC  YLHPDLYERGVLRKHGNKAFSATVDLWSIGVTFYHIATGQLPFRPYGG-RQNKVTMYEIT
IKKe_HUMAN        YLHPDMYERAVLRKPQQKAFGVTVDLWSIGVTLYHAATGSLPFIPFGGPRRNKEIMYRIT
TBK1_HUMAN        YLHPDMYERAVLRKDHQKKYGATVDLWSIGVTFYHAATGSLPFRPFEGPRRNKEVMYKII
                  *****:***.****  :* :.. *********:** ***.*** *: * *.*.  *: *

IKKeTBK1_APOC     SKKRSGMISGVQKSEGEDIEWSDKLPEHTRLSQGLKDLLTPVLAGILESEPSNA-MSFEE
IKKeTBK1_AMIL_v3  SKKKPGVISGVQKSEGEEIEWSDKLPEHTRLSQGLKVLFTPVLAGALESEPCSTSMSFEQ
IKKeTBK1_AMIL_v6  SKKKPGVISGVQKSEGEEIEWSDKLPEHTRLSQGLKVLFTPVLAGALESEPCSTSMSFEQ
IKKeTBK1_ACER_BC  SKKKPGIISGVQKSEGEEIEWSDKLPENTRLSQGLKVLFTPVLAGALESEPCSTSMSFEQ
IKKeTBK1_ACER_CC  SKKKPGIISGVQKSEGEEIEWSDKLPENTRLSQGLKVLFTPVLAGALESEPCSTSMSFEQ
IKKE_HUMAN        TEKPAGAIAGAQRRENGPLEWSYTLPITCQLSLGLQSQLVPILANILEVEQAKC-WGFDQ
TBK1_HUMAN        TGKPSGAISGVQKAENGPIDWSGDMPVSCSLSRGLQVLLTPVLANILEADQEKC-WGFDQ
                  : * .* *:*.*. *.  ::**  :*    ** **:  :.*:**. ** :  .   .*::

IKKeTBK1_APOC     FFARIQDILSRKVIDVYSVHSASFHKIYIKQDETFAKFQELIAVQTGVSAAQQKLFFKYD
IKKeTBK1_AMIL_v3  FFASVQDILTRKVIDIYSVHSACFHKIYMKPNETFAKFQELIAVQTGVSARQQMLMFKYE
IKKeTBK1_AMIL_v6  FFASVQDILTRKVIDIYSVHSACFHKIYMKPNETFAKFQELIAVQTGVSARQQMLMFKYE
IKKeTBK1_ACER_BC  FFTSVQDILTRKVIDIYSVHSACFHKIYMKPNETFAKFQELIAVQTGVSARQQMLMFKYE
IKKeTBK1_ACER_CC  FFTSVQDILTRKVIDIYSVHSACFHKIHMKPNETFAKFQELIAVQTGVSARQQMLMFKYE
IKKE_HUMAN        FFAETSDILQRVVVHVFSLSQAVLHHIYIHAHNTIAIFQEAVHKQTSVAPRHQEYLFEGH
TBK1_HUMAN        FFAETSDILHRMVIHVFSLQQMTAHKIYIHSYNTATIFHELVYKQTKIISSNQELIYEGR
                  **:  .*** * *: ::*: .   *:*:::  :* : *:* :  ** : . :*  :::
```

**Supplemental Figure 13.** Alignment of coral and human IKKE/TBK1 proteins (continued)

```
IKKeTBK1_APOC     EFRPDPMAASSSYPNTSDDNLMVMVGGEYMSPERLFLHKIPKL--PKMPEE-YTCEADSA
IKKeTBK1_AMIL_v3  EFRPDPMAPASSYPNTTEDSFMVMIGGDSILPDRLNFFKIPKM--PNVPEEGTDRESDAS
IKKeTBK1_AMIL_v6  EFRPDPMAPASSYPNTTEDSFMVMIGGDSILPDRLNFFKIPKM--PNVPEEGTDRESDAS
IKKeTBK1_ACER_BC  EFRPDPMAPASSYPNTTEDSFMVMIGGDSILPDRLNFFKIPKM--PNVPEEGTDLESDAS
IKKeTBK1_ACER_CC  EFRPDPMAPASSYPNTTEDSFMVMIGGDSILPDRLNFFKIPKM--PNVPEEGTDLESDAS
IKKE_HUMAN        LCVLEPSVSAQHIAHTTASSPLTLFSTA--IPKGL-AFRDPALDVPKFVPK-VDLQADYN
TBK1_HUMAN        RLVLEPGRLAQHFPKTTEENPIFVVSREPLNTIGLIYEKISL---PKVHPR-YDLDGDAS
                      :*   :.  .:*: .. : :..     .  *   . .    *:.       :.*

IKKeTBK1_APOC     IAKMMANRAYCCKYAVQQYFRATKAMLVTIESILKNLKKDMMLYVSFCKQIESDWTALQQ
IKKeTBK1_AMIL_v3  VAKIMTTRAHYCKYAVELYTRATKAMLLTVESVIKNLKKVMMLYISSCKQVESQSIALNQ
IKKeTBK1_AMIL_v6  VAKIMTTRAHYCKYAVELYTRATKAMLLTVESVIKNLKKVMMLYISSCKQVESQSIALNQ
IKKeTBK1_ACER_BC  VAKIMTTRAHYCKYAVELYTRATKAMLLTVESVIKNLKKVMMLYISSCKQVESQSIALNQ
IKKeTBK1_ACER_CC  VAKIMTTRAHYCKYAVELYTRATKAMLLTVESVIKNLKKVMMLYISSCKQVESQSIALNQ
IKKE_HUMAN        TAKGVLGAGYQA-------LRLARALLDGQELMFRGLHWVMEVLQATCRRTLEVARTSLL
TBK1_HUMAN        MAKAITGVVCYA-------CRIASTLLLYQELMRKGIRWLIELIKDDYNETVHKKTEVVI
                   ** :      .        * : ::*   * : ..:.. : :     .

IKKeTBK1_APOC     VSEVSCNSQSHLIWVFEALLSKITEPQ-DELQLIWDEVQDIKKWTDSKRQSEAELRKTNK
IKKeTBK1_AMIL_v3  TCDVCCSSHDHLVGIFDALVSQFTEPQ-DKIESLQEKIQTIKKWTDAKRQSEKELHKINE
IKKeTBK1_AMIL_v6  TCDVCCSSHDHLVGIFDALVSQFTEPQ-DKIESLQEKIQTIKKWTDAKRQSEKELHKINE
IKKeTBK1_ACER_BC  TCDVCCSSHDHLVRVFDTLVSQFTEPQ-DKIESLQEKIQTIKKWTDAKRQSEKELHKINE
IKKeTBK1_ACER_CC  TCDVCCSSHDHLVRVFDTLVSQFTEPQ-EKIESLQEKIQTIKKWTDAKRQSEKELHKINE
IKKE_HUMAN        YLSSSLGT--------ERFSSVAGTPEIQELKAAAELRSRLRTLAEVLSRCSQNITETQE
TBK1_HUMAN        TLDFCIRNIEKTVKVYEKLMKINLEAA--ELGEISDIHTKLLRLSSSQGTIETSLQDIDS
                    . .  .        : : .    .   ::    :    :   :.     . .: . :.

IKKeTBK1_APOC     GLDRFRSLISDVIETDRFYTFLETFRKDAEDPYLFNVVSTYATNIEEIYQSFRKDKHHTS
IKKeTBK1_AMIL_v3  GLQQFRDLISSVVEKDRFLPFLEEFRSDTEKEKFFNLVSTYTRNTADIYTLFRKDKHQ--
IKKeTBK1_AMIL_v6  GLQQFRDLISSVVEKDRFLPFLEEFRSDTEKEKFFNLVSTYTRNTADIYTLFRKDKHQ--
IKKeTBK1_ACER_BC  GLQQFRDLISSVVEKDRFLPFLEEFRSDTEKEKFFNLVSTYTRNTADICTLFRKDKNQ--
IKKeTBK1_ACER_CC  GLQQFRDLISSVVEKDRFLPFLEEFRSDTEKEKFFNLVSTYTRNTADICTLFRKDKNQ--
IKKE_HUMAN        SLS---SLNRELVKS----------RDQVHEDRSIQQIQCCLDKMNFIYKQFKKSRMR--
TBK1_HUMAN        RLSPGGSLADAWAHQ----------EGTHPKDRNVEKLQVLLNCMTEIYYQFKKDKAE--
                   *.   .*                      .   .: :.        *   *.*..

IKKeTBK1_APOC     QNRRLSYADEQRHIFDRKKIVSLCNK-ITEVTDESLGKRSELHGQLVQWLNEVNRCYDEA
IKKeTBK1_AMIL_v3  --RRLTVADEQRHAFDRKKIVVQCEK-ITRVTDDALGQRSQLHAKLIHWLSEVDACTEEA
IKKeTBK1_AMIL_v6  --RRLTVADEQRHAFDRKKIVVQCEK-ITRVTDDALGQRSQLHAKLIHWLSEVDACTEEA
IKKeTBK1_ACER_BC  --RRLTVADKQRHAFDRKKIVVQCEK-ITRVTDDALGQRSQLHAKLIHWLSEVDTCTEEA
IKKeTBK1_ACER_CC  --RRLTVADKQRHAFDRKKIVVQCEK-ITRVTDDALGQRSQLHAKLIHWLSEVDTCTEEA
IKKE_HUMAN        --PGLGYNEEQIHKLDKVNFSHLAKRLLQVFQEECVQKYQASLVTHGKRMRVVHETRNHL
TBK1_HUMAN        --RRLAYNEEQIHKFDKQKLYYHATKAMTHFTDECVKKYEAFLNKSEEWIRKMLHLRKQL
                      *   ::* * :*. ::   . . :  . ::.: : .        .:  :    .

IKKeTBK1_APOC     ENFHNTFVGQTQAKDAYLFALQTVQKECKEKSKKIVQKLQKV--ITSSSSVPERTEPGPI
IKKeTBK1_AMIL_v3  ENLQNSFVVHTEAMEAHRLSLQTLQEQCQVKSKEILQEIQQLQSLTTSNAASHPGSSSPV
IKKeTBK1_AMIL_v6  ENLQNSFVVHTEAMEAHRLSLQTLQEQCQVKSKEILQEIQQLQSLTTSNAASHPGSSSPV
IKKeTBK1_ACER_BC  ENLQNSFVVHTEAMEAHRISLQTLQEQCQVKSKEILQEIQQLQSLTTSNVASHPGSSSPV
IKKeTBK1_ACER_CC  ENLQNSFVVHTEAMEAHRISLQTLQEQCQVKSKEILQEIQQLQSLTTSNVASHPGSSSPV
IKKE_HUMAN        RLVGCS-----------VAACNTEAQGVQESLSKLLEELSHQ--LLQDRAKGAQASPPPI
TBK1_HUMAN        LSLTNQ-----------CFDIEEEVSKYQEYTNELQETLPQK--MFTA-SSGIKHTMTPI
       .           :  .  :   .:: : : :  :          *:
```

**Supplemental Figure 13.** Alignment of coral and human IKKE/TBK1 proteins (continued)

```
IKKeTBK1_APOC     TENGHTDP--VINGVTMNQLGNELDSLSVITKESVRTAKQNTQQIESVIGSFIKLYSSIN
IKKeTBK1_AMIL_v3  L-NVHEEQEEQILTDTMQALGYELDTLSHITTASLGTAKDNTAQIERLVSGLIKSYESMN
IKKeTBK1_AMIL_v6  L-NVHEEQEEQILTDTMQALGYELDTLSHITTASLGTAKDNTAQIERLVSGLIKSYESMN
IKKeTBK1_ACER_BC  L-NGHEEQEEQILTDTMQALGYELDTLSDITTASLGTAKDNTAQIERLISGLIKSYESMN
IKKeTBK1_ACER_CC  L-NGHEEQEEQILTDTMQALGYELDTLSDITTASLGTAKDNTAQIERLISGLIKSYESMN
IKKE_HUMAN        A---------PYPSPTRKDLLLHMQELC--------------EGMKLLASDLLD-----N
TBK1_HUMAN        Y---------P-SSNTLVEMTLGMKKLK--------------EEMEGVVKELAE-----N
                                 *   :   :. *                 :: :   : .     *

IKKeTBK1_APOC     EHLMKNVEQLHETVAEESQN--
IKKeTBK1_AMIL_v3  DHLTKNFEQLVEAVRGEADHLQ
IKKeTBK1_AMIL_v6  DHLTKNFEH-------------
IKKeTBK1_ACER_BC  DHLKKNFEQ-------------
IKKeTBK1_ACER_CC  DHLKKNFEQLVEAVRGEADHLQ
IKKE_HUMAN        NRIIERLNRVPAPPDV------
TBK1_HUMAN        NHILERFGSLTMDGGLRNVDCL
                  :.: :..
```

Blackbird & Calabash IKKE/TBK1 protein sequences (MZ812724, MZ812725) versus human, *Acropora millepora* (AMIL) and *Astrangia poculata* (APOC; [8]). The Blackbird sequences lacks 13 residues at the carboxy terminus that are found in the Calabash sequence. Similarly, one of the *A. millepora* sequences (version 6) lacks the same 13 residues found at the carboxy terminus of the other *A. milleopora* sequences (version 3). The sequences were aligned using the program MUSCLE (version 3.8.31; [1]) as implemented on the Phylogeny.fr web server [2]. Asterisks denote positions that are strictly conserved. Colons and periods denote conservative substitutions.

### **Supplemental Figure 14.** Conserved motifs in coral and human IKK proteins

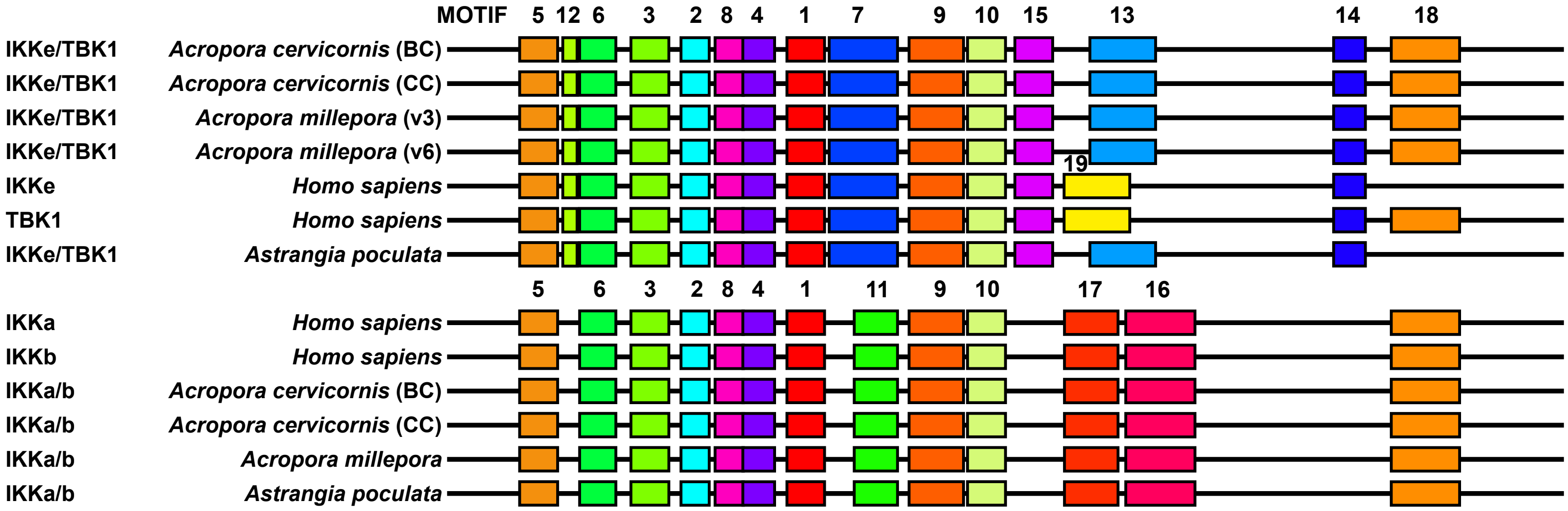


**Supplemental Figure 14.** Diagrammatic representation of IKK proteins in corals and humans showing the location of conserved motifs. The motifs were originally identified by Nguyen in a phylogenetically diverse collection of metazoan IKK proteins using MEME [9,10]. These previously defined motifs were located in *Acropora* IKK proteins in the current study using MAST [11]. Motifs are numbered in order of their statistical significance, from most significant (lowest E-value) to least significant. By definition, the length of a given motif, represented by the width of each colored box, is conserved in all instances of that motif. Intermotif distances (represented by the thin black line) are not drawn to scale.

**Supplemental Figure 15.** Distance tree relating coral colonies based on 35,637 polymorphic nucleotide sites

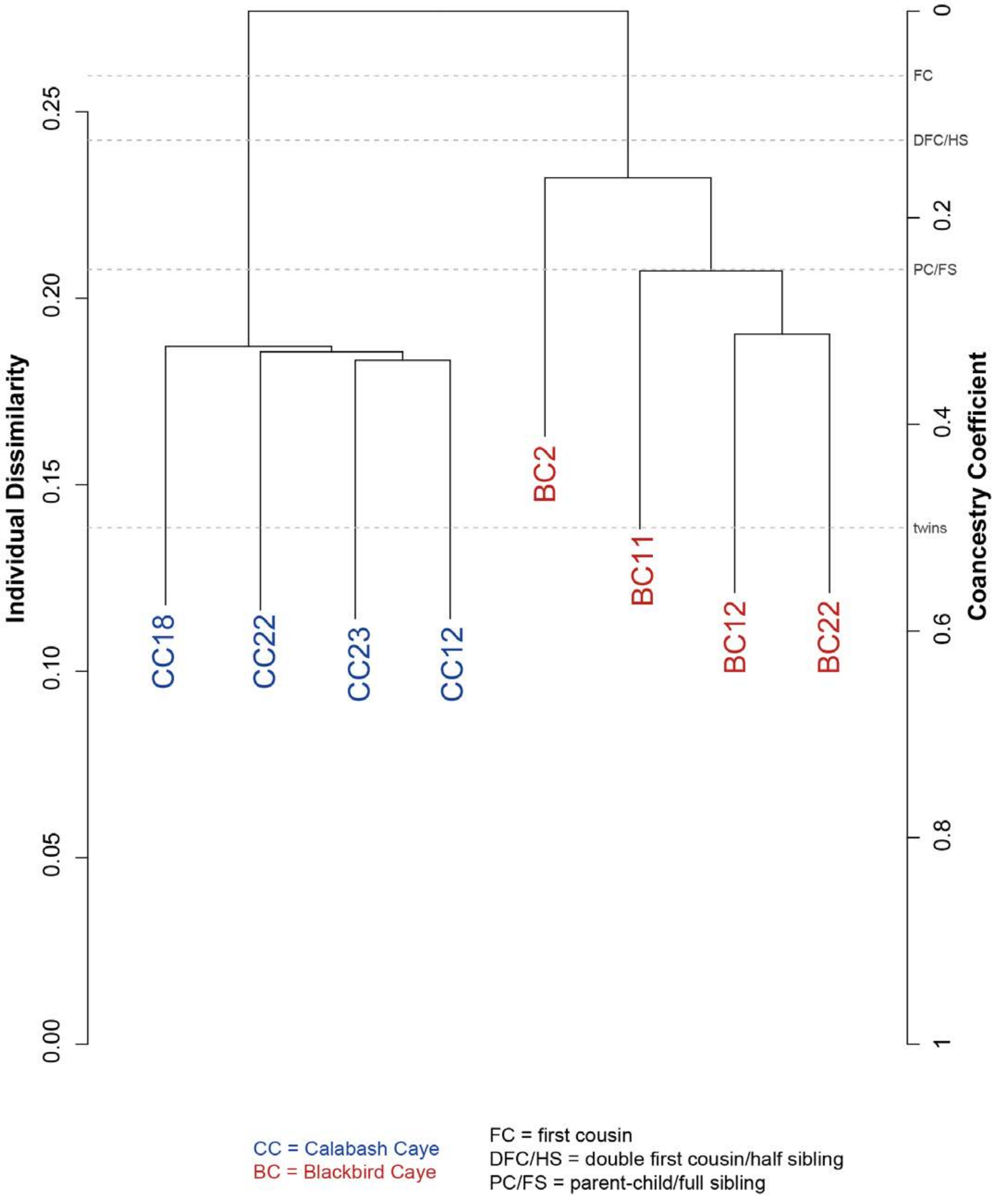

**Supplemental Figure 15.** Distance tree relating coral colonies based on 35,637 polymorphic nucleotide sites. To characterize coral genetic diversity within and between sites, we identified SNPs (single nucleotide polymorphisms) as follows: All sequenced libraries were mapped to the concatenated holobionts transcriptome using bowtie2 (Langmead and Salzberg, 2012). SNP calling was performed on the processed SAM files using SAMtools (Li et al., 2009). SNPs that mapped to contigs in the taxclasses “Cnidaria” and were filtered using VCFtools (Danecek et al., 2011). We retained only those SNPs meeting the following criteria: (1) genotype and site quality scores > 30; (2) minimum read depth ≥ 2X the mean read depth for SNPs in the Cnidaria contig bin (≥170); (3) minor allele frequency ≥0.05; (4) no missing values, which resulted in 35,637 SNPs. We used the packages SNPRelate v. 1.20.1 (Zheng et al., 2012) to perform a hierarchical clustering analysis based on the calculated identity-by-state values. The snpgdsDrawTree function in this package plots the dendrogram and thresholds for estimated relationships among individuals (e.g., twins, parent/child, half sibling).

**Supplemental Figure 16.** Example calculations for taxonomic, CPM, and gene purity.

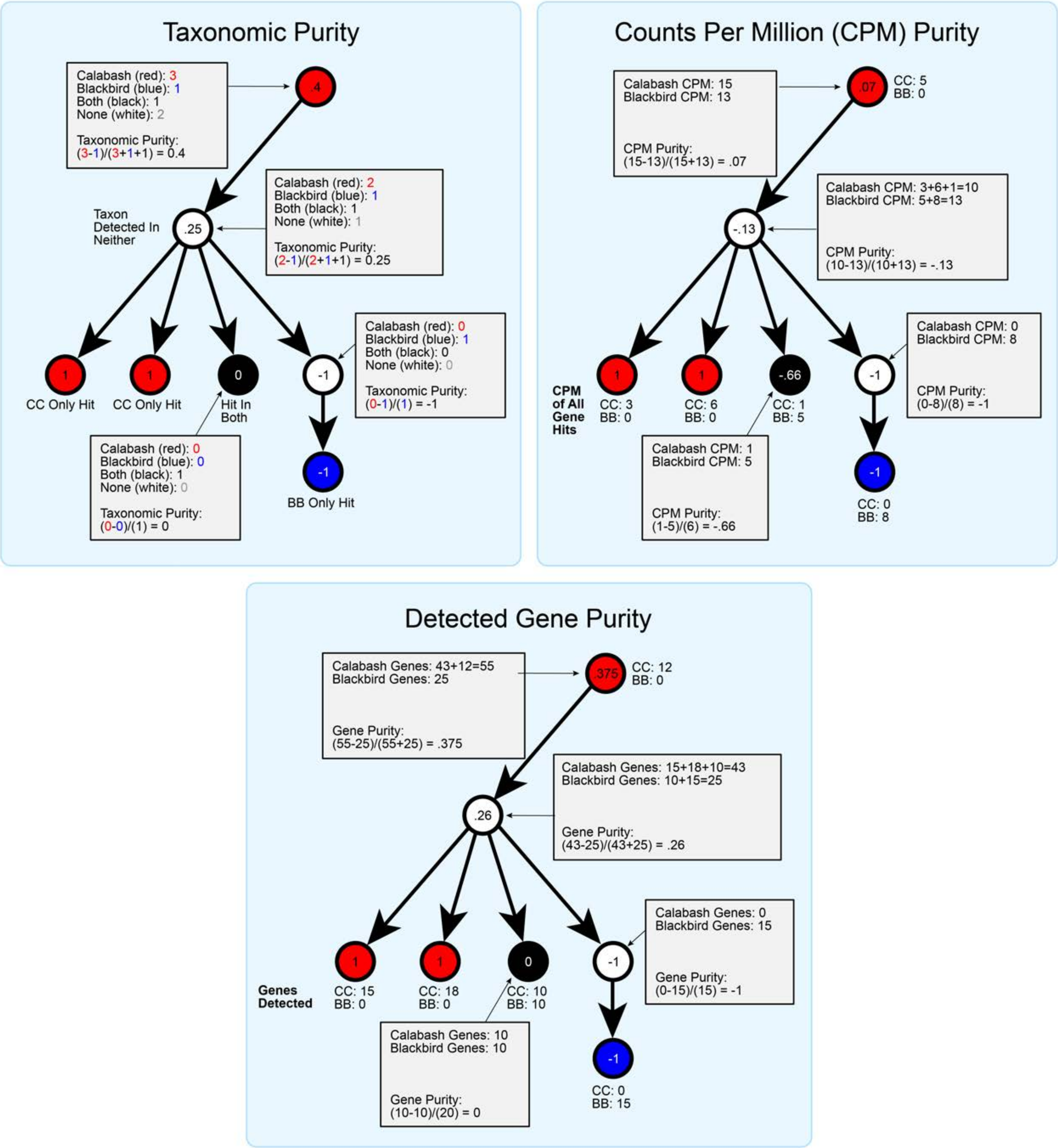

**Supplemental Figure 16.** Example calculations for taxonomic, CPM, and gene purity.

All three purity score calculations use the “directly detected” nodes at and below each node to compute purity. Taxonomic purity is calculated by counting the number of nodes that were directly detected in Calabash, Blackbird, or both at each node and below. For leaf nodes detected in only one site, the purity score is either 1 or -1. For ancestor nodes, the purity is calculated as the difference in the number of Calabash vs Blackbird exclusive taxa, divided by the total number of directly detected taxa at and below each node. Gene purity is calculated similarly, except now the number of genes/protein sequences with a hit from each site are counted, rather than the simple presence/absence used in taxonomic purity. Both taxonomic and gene purity are assessed using only the assembled transcripts from each site. CPM purity is slightly different than the other two in that the RNASeq data used in the assembly is instead used to quantify the abundance of hits in each taxon. After summing the CPM mapping to each taxon from each site, the purity score is calculated similarly by dividing the difference in CPM mapping from each site by the total CPM detected across both sites.

**Supplemental Figure 17.** Site-specific purity scores for taxa, genes and normalized RNA read counts.

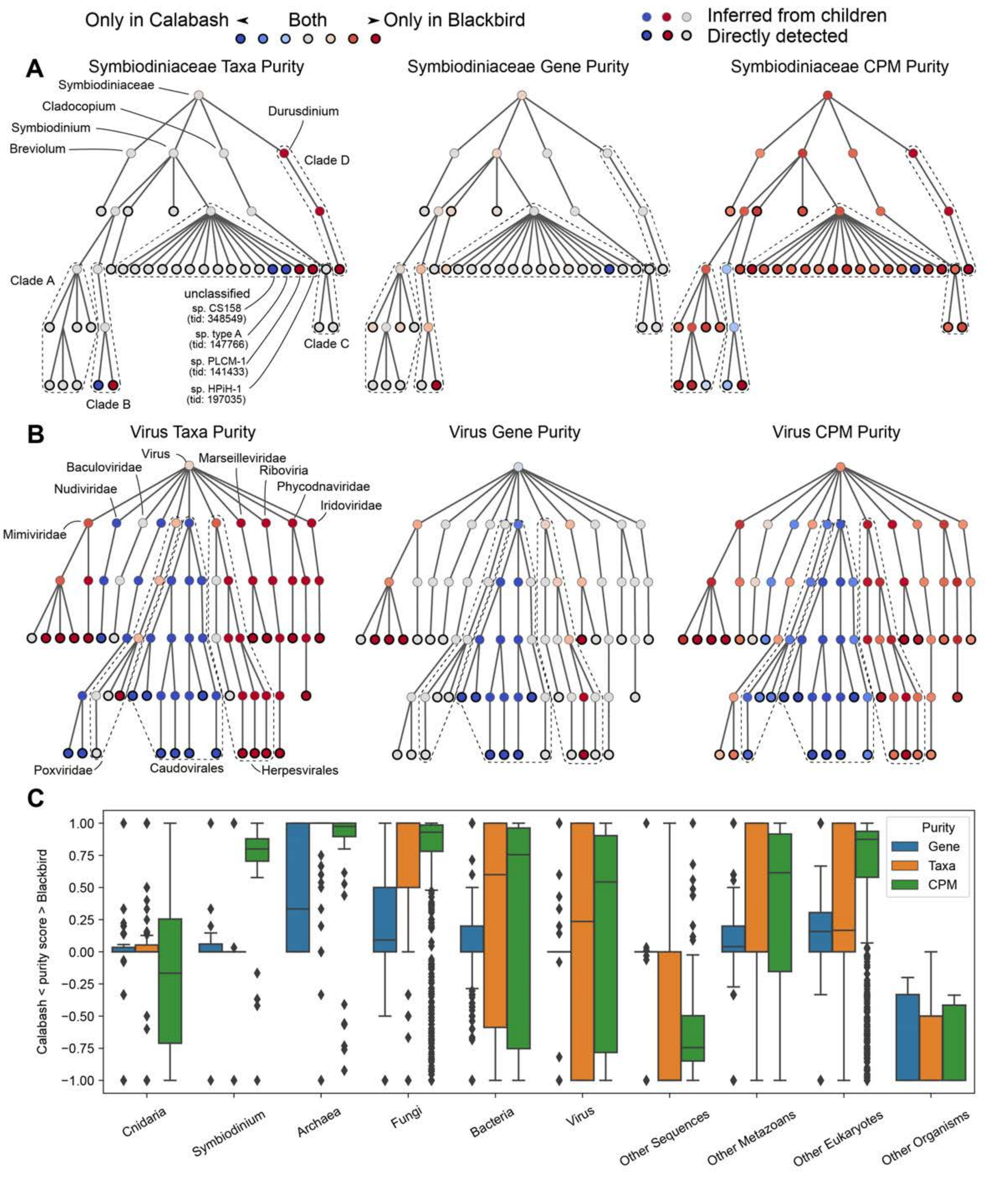

**Supplemental Figure 16.** Graphs depicting the degree of taxonomic skewness between sites for **(A)** Symbiodiniaceae and **(B)** viruses. In each graph, taxa are represented by nodes, which are ordered by increasing taxonomic specificity from top to bottom. The color of each node represents its purity score, with blue nodes biased toward Calabash and red nodes biased toward Blackbird.**(C)** Distribution boxplots of purity scores for directly detected taxa across all taxclasses. Box bodies and whiskers indicate inner quartile range and 5$^{th}$/95$^{th}$ percentile of corresponding scores, respectively. Narrow boxes near zero indicate no skew between sites; tall boxes indicate a broad diversity of site-specificity; boxes near 1 or -1 indicate strong skew to Calabash or Blackbird, respectively. Detailed information is provided below.

The purity scores describe the skew of representation between Blackbird and Calabash sites with respect to the taxa, genes, and normalized RNASeq read counts (counts per million or CPM) across the entire taxonomic tree. The taxonomic tree is represented as a directed graph where nodes are individual taxa and directed edges indicate taxonomic parent-child relationships. Every taxon in the tree has a path to the "root" taxon, which is an abstract taxon representing all forms of life. Walking down from this root taxon through the tree visits nodes (taxa) of increasing specificity, e.g. "All life" -> cellular organisms -> Bacteria -> Proteobacteria -> Gammaproteobacteria -> Enterobacteriales -> Enterobacteriaceae -> Escherichia -> Escherichia coli. Each of these intermediate taxa have their own taxonomic ID in the NCBI taxonomic database. Note the traditional Linnean taxonomic levels e.g. species are not strictly represented in the structure of the tree but are annotated to nodes as appropriate. This tree structure enables information on more specific taxa to be summarized to more general taxa at every node in the taxonomic tree. The purity score algorithm takes advantage of this structure to compute purity for each node using attributes of all its child nodes. In this way, entire subtrees of the taxonomy can be easily identified as being skewed to one site or the other without being biased by aspects of the structure of the tree, e.g. taxa with different numbers of subtaxa, due to uneven representation of annotated taxa in the database. For example, there are many more strains of *E. coli* annotated in the database than Cnidaria, and the results should not be influenced by these imbalances.

The taxonomic tree structure described above also enables partitioning the taxonomy into disjoint subtrees representing different groups of organisms. Given a set of taxon IDs within the tree, a lowest common ancestor algorithm may be employed to identify the largest subtree within the taxonomy that contains all those taxon IDs. For example, one might provide the taxon IDs for all Cnidaria species to identify the smallest subtree that contains all of these species. This subtree can then be removed from the larger taxonomy, leaving the species specific tree and the remaining taxa as disjoint trees. This is the strategy employed by the taxonomic parsing algorithm used to separate transcripts into taxclasses, i.e. Cnidaria, Symbiodinium, Archaea, Fungi, Bacteria, Virus, Other Metazoans, Other Eukaryotes, Other Organisms, and Other Sequences (NCBI taxid 28384).

The taxonomic tree for each taxclass is hereafter called a "taxgraph". Each NCBI protein or nucleotide sequence is annotated to one or more NCBI taxids. A sequence may be annotated to more than one taxid if the exact sequence is shared by multiple organisms. We therefore can annotate the nodes in each taxgraph with their genes/proteins and also record which site had transcripts or reads that map to these sequences. When a node annotated with a gene/protein also has a matching transcript from either site as determined by minimap2 or DIAMOND, that node is marked as "directly detected." Nodes that have no annotated sequences, or that do but lack any direct match to a transcript, are not directly detected but are still useful as intermediate or "inferred" taxa within the graph. Each taxgraph is pruned such that all leaf nodes (i.e. nodes without any children) are directly detected, since more specific taxa in the taxgraph that are not detected are not informative to the purity algorithm.

The full taxgraph_stats.csv file can be downloaded from: https://bitbucket.org/adamlabadorf/staghorn_transcriptome/src/master/analysis/taxgraph_stats.csv

This csv can easily be sorted and filtered for browsing and exploration in Excel. As the proportion of Bacteria taxclass transcripts differed greatly between sites, we decided to further examine the Bacteria superkingdom purity results. To do so, we first filtered the "superkingdom" column to only display "Bacteria" rows, and then filtered the "rank" column to only display the "phylum" level entries. Next, sorting the "taxa_diff" column by descending values allowed us to focus on phyla that many have lower taxonomic classes weighted towards presence at one site. Scanning the first several rows under the column "taxa_purity", we see the highest value, 0.933 (which is also the largest absolute value), corresponds to Planctomycetes. This score indicates that members of this phylum are found almost exclusively at Blackbird, as it is close to +1. The "cpm_purity" and "gene_purity" values are 0.934 and 0.289 respectively.

**Supplemental Table 1.** Taxonomic class definitions and properties

| Priority* | taxclass (description) | Taxon ID |
|---|---|---|
| 0 | Other Organisms (any taxon IDs not belonging to taxclasses 1-9) | NA |
| 1 | Bacteria | 2 |
| 2 | Fungi | 4751 |
| 3 | Archaea | 2157 |
| 4 | Viruses | 10239 |
| 5 | Other Sequences (BACs, transposons, etc.) | 28384 |
| 6 | Other Eukaryotes (eukaryotes except taxclasses 2, 7, 8, or 9) | 2759 |
| 7 | Other Metazoa (metazoans other than Cnidaria) | 33208 |
| 8 | Cnidaria | 6073 |
| 9 | Symbiodiniaceae | 252141 |
| 10 | No Hit (sequences that have no BLAST hits) | NA |

* Priority column indicates how BLAST hits mapping to multiple taxclasses are resolved, where smaller priority numbers are preferred over higher.

**Supplemental Table 2.** Recovery of NF-kB signaling proteins from Blackbird and Calabash transcriptomes.

| Gene | Loc (version)[1] | Complete or Partial[2] | Length[3] | Identity BC vs CC | Accession |
|---|---|---|---|---|---|
| **NF-kB** | Blackbird | Complete | 904 aa | 904/904 | *MZ851284* |
| | Calabash | Complete | 904 aa | | *MZ501696* |
| **IKK a/b** | Blackbird | Complete | 731 aa | 731/731 | *MZ812722* |
| | Calabash | Partial | 731 aa | | *MZ812723* |
| **IKKe/TBK1** | Blackbird | Complete | 779 aa | 779/792 | *MZ812724* |
| | Calabash | Partial | 792 nt | | *MZ812725* |

**Supplemental Table 3.** Recovery of Symbiodiniaceae molecular markers from Blackbird and Calabash.

| Gene | Loc (version)[1] | Complete or Partial[2] | Length[3] | Identity BC vs CC | Phylogenetic affinity within Symbiodiniaceae | Accession |
|---|---|---|---|---|---|---|
| **Actin** | Blackbird (1) | Complete | 376 aa | 376/376 | Clade A / *Symbiodinium* | *MZ501686* |
| | Calabash (1) | Complete | 376 aa | | Clade A / *Symbiodinium* | *MZ501688* |
| | Blackbird (2) | Complete | 375 aa | 356/375 | Clade A / *Symbiodinium* | *MZ501687* |
| | Calabash (2) | Complete | 375 aa | | Clade A / *Symbiodinium* | *MZ501689* |
| **COI** | Blackbird | Complete | 491 aa | 442/454 | Clade A / *Symbiodinium* | *MZ501690* |
| | Calabash | Partial | 454 aa | | Clade A / *Symbiodinium* | *MZ501691* |
| **ITS-2** | Blackbird | Complete | 251 nt | 118/118 | Clade A / *Symbiodinium* | *MZ503278* |
| | Calabash | Partial | 118 nt | | Clade A / *Symbiodinium* | *NA** |
| **Rad24** | Blackbird (1) | Complete | 240 aa | 214/214 (1)<br>204/214 (2) | Clade A / *Symbiodinium* | *MZ501692* |
| | Blackbird (2) | Complete | 241 aa | | Clade A / *Symbiodinium* | *MZ501693* |
| | Calabash | Complete | 214 aa | | Clade A / *Symbiodinium* | *MZ501694* |

* >ACER_Calabash ITS2 partial
TTTTTATCAGTCACAACAGCAAGCGAAACTTGGTACTTAGCATGCCAGTGCCAATTGCAGAAGCATGCA
GCAGCACTGCTCCTGATAACAAGAGCAGCAGAAAAGAAAGTAGAAGCACAAGATCGGAAGAGCACA

**Supplemental Table 4.** Number of transcripts shared by and unique to Blackbird and Calabash by taxclass

| Taxclass | Total Transcripts | % of Total | Both sites | % of Taxclass | Blackbird only | % of Taxclass | Calabash only | % of Taxclass |
|---|---|---|---|---|---|---|---|---|
| **Cnidaria** | 684596 | 68.3% | 659357 | 96.3% | 18730 | 2.7% | 6509 | 1.0% |
| **Symbiodiniaceae** | 85605 | 8.5% | 32680 | 38.2% | 52828 | 61.7% | 97 | 0.1% |
| **Bacteria** | 36639 | 3.7% | 12016 | 32.8% | 1939 | 5.3% | 22684 | 61.9% |
| **Other Metazoa** | 26898 | 2.7% | 17019 | 63.3% | 7182 | 26.7% | 2697 | 10.0% |
| **Other Eukaryota** | 11740 | 1.2% | 5757 | 49.0% | 5293 | 45.1% | 690 | 5.9% |
| **Fungi** | 1431 | 0.1% | 689 | 48.1% | 628 | 43.9% | 114 | 8.0% |
| **Other Organisms** | 1114 | 0.1% | 502 | 45.1% | 328 | 29.4% | 284 | 25.5% |
| **Other Sequences** | 858 | 0.1% | 518 | 60.4% | 205 | 23.9% | 135 | 15.7% |
| **Viruses** | 225 | 0.0% | 99 | 44.0% | 61 | 27.1% | 65 | 28.9% |
| **Archaea** | 97 | 0.0% | 36 | 37.1% | 31 | 32.0% | 30 | 30.9% |
| **No Hit** | 153086 | 15.3% | 95668 | 62.5% | 39492 | 25.8% | 17926 | 11.7% |
| **All Taxclasses** | 1002289 | 100.0% | 824341 | 82.2% | 126717 | 12.6% | 51231 | 5.1% |
| **Non-Cnidaria** | 317693 | 31.7% | 164984 | 51.9% | 107987 | 34.0% | 44722 | 14.1% |
